# The cerebellum exploits a thalamic carrier frequency to refine motor behaviors

**DOI:** 10.64898/2026.09.13.751058

**Authors:** Brielle Miano, Zhiyi Yao, Elise Dailey, Zekiel Factor, Jonathan E Ploski, Christopher H. Chen

## Abstract

Despite enabling nearly all cerebellar-cortical communication, how the thalamus processes and interacts with cerebellar activity remains unclear. We found a pronounced 1-5 Hz (*delta)* rhythm in the thalamic membrane potential and cerebellothalamocortical local field potentials during nonmoving periods in behaving mice. We show, using photometry, multielectrode recordings, closed-loop stimulation, and dynamic clamp, that CaV3.1-dependent resonance of thalamic neurons facilitates transmission of *delta-*rhythm cerebellar activity to the cortex. Knockout of CaV3.1 in cerebellar-thalamic neurons diminishes the thalamocortical *delta* oscillation and specifically impairs motor learning while minimally affecting motor performance. These results ground population oscillations to features of ion channels, highlight the cerebellum and *delta*’s utility in offline motor learning, and support a carrier frequency-like encoding mechanism that underlies cerebellothalamocortical communication.

## MAIN TEXT

High-fidelity communication across brain regions is challenging, requiring specific adaptations to overcome many potential points of failure across multisynaptic pathways. Binding communication to local oscillations (communication through coherence)^1–5^ is an attractive framework that has been used extensively to explain the role of *gamma* and *theta* rhythms across the cortex and hippocampus.

This might also be accomplished by the resonant properties of thalamic neurons. CaV3.1-mediated low-threshold spikes (LTS) amplify subthreshold inputs and boost the thalamic response to specific input frequencies (1-5 Hz)^6,7^. CaV3.1 dependence also means that resonance within this frequency band is dependent on thalamic membrane potential. Deinactivation of CaV3.1 at hyperpolarized

potentials puts the thalamus in “burst mode,” which features nonlinear LTS burst firing interspersed with biophysically constrained pauses in time with resonant frequencies^8^. At more depolarized potentials thalamic cells enter “tonic mode,” where regular spiking dominates. Because tonic firing is thought to be much more permissive for information transmission, burst mode is often implicated in attention-gating behavioral states^8–11^, although its precise functional role remains an open question.

The cerebellothalamocortical motor pathway is a critically important, rapidly regulated, and multisynaptic route that might exploit thalamic resonance. Cerebellar efferents have uniquely high baseline activity: each deep cerebellar nuclei (DCN) neuron spontaneously fires at rates up to 50 spikes/s even during motor quiescence^12–14^, and single motor thalamic neurons receive multiple DCN inputs^15^. Rhythmic population-level activity is encoded in the cerebellum^16–19^. Consequently, if aggregate population-level activity of the DCN oscillated in-phase with resonating thalamic activity, cerebellar activity would be propagated to the cortex even during the “gated” thalamic burst mode.

## RESULTS

### The cerebellum, thalamus and cortex share *delta*-band oscillatory activity during stillness

Since trans-thalamic communication must rely on firing mode, we first investigated how firing mode relates to behavioral state. We performed whole-cell *in vivo* patch-clamp in cerebellar-recipient thalamic territories in awake mice through periods of movement and stillness (**Figure 1A-H**). We identified these territories by using the ChR2-evoked field-response from mice injected with AAV1-ChR2 in the DCN (**Figure 1A, B**) and found that motor thalamic cells exhibit tonic spiking during movement (**Figure 1D**). During periods of stillness when animals are alert and maintaining a fixed body position, thalamic cells enter burst mode, as evidenced by LTS events (**Figure 1E**), hyperpolarized membrane potentials (**Figure 1F**), and wider membrane potential variance reflecting up- and down-states (**Figure 1G**). Membrane potentials also show enhanced *delta* oscillatory activity selectively in still periods (**Figure 1D, H**), demonstrating a functional link between thalamic burst mode and *delta* oscillatory activity during stillness.

**Figure 1.**
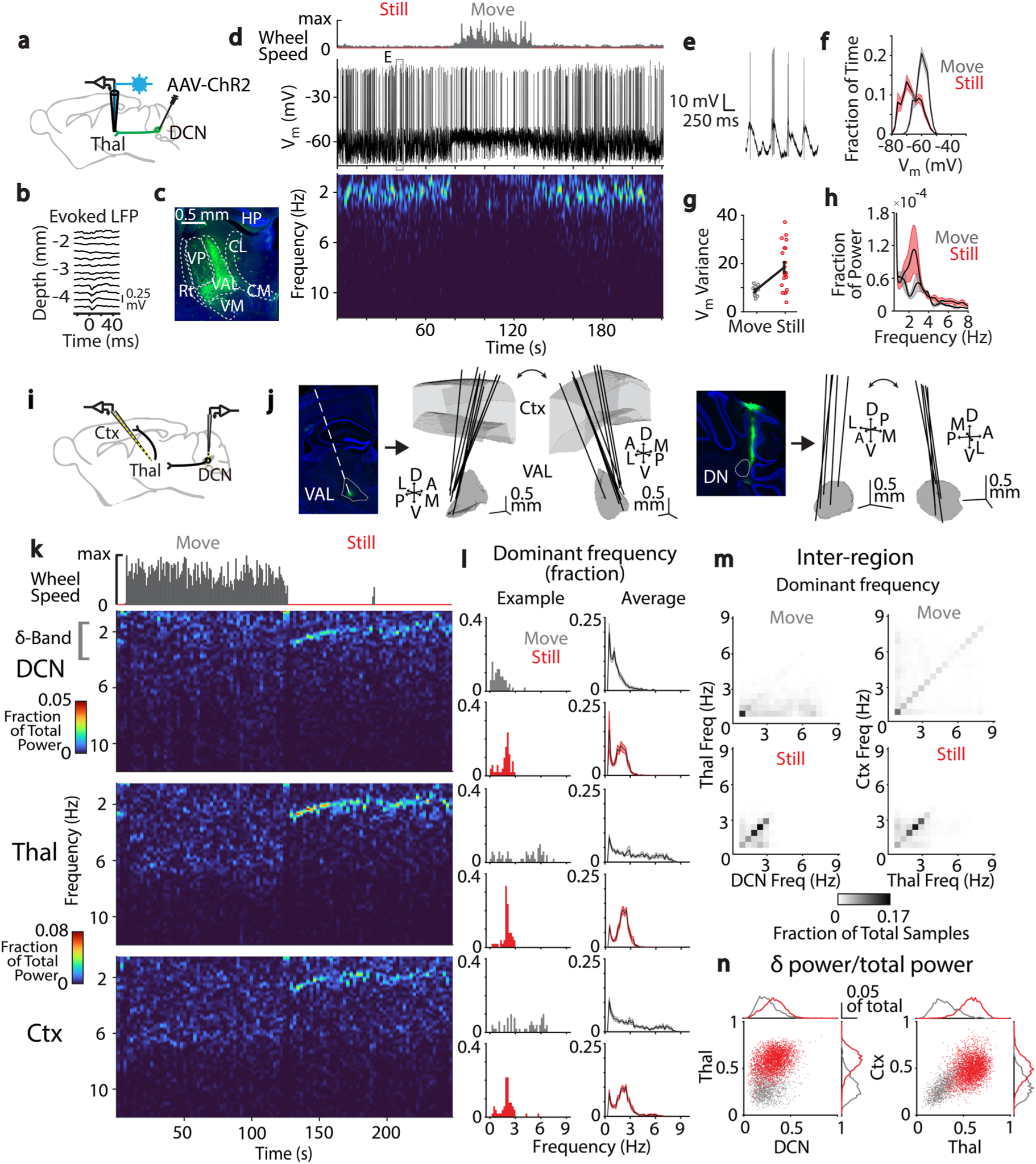
Coherent *delta*-band oscillatory activity in DCN, thalamus, and cortex. **(A-H)** AAV1-ChR2-YFP was injected into the DCN, and awake, *in vivo* whole-cell recordings were made in the thalamus. **(B)** Light pulses (10 ms) evoking depth-dependent field responses were used to identify cerebellar motor thalamus. **(C)** Coronal section of thalamus confirming ChR2+ fibers (green) from DCN. **(D)** Example trace from whole-cell recording in thalamus. **(E)** Indicated recording segment in (**D**) showing plateau potentials and burst firing. **(F)** Average membrane potential distributions for still/moving episodes (n = 4 cells). **(G)** Membrane potential variability in still vs. move conditions for different still/move epochs. **(H)** Power spectrum of membrane potential in each movement condition. **(I-N)** Triple-site silicon probe electrophysiological recordings in cortex, motor thalamus and DCN were performed. **(J)** Left: Sample recording tract through cortex and thalamus (green: tip of probe in ventral anterior-lateral thalamus, VAL) with locations of all cortex-thalamus probe tracts (n = 9 insertions, N = 5 animals). Right: same as left, for DCN probe tracts (n = 5 insertions, N = 3 animals). Trajectories plotted on normalized mouse brain. DN: dentate nucleus of the DCN. **(K)** Example spectrograms of simultaneous extracellular recordings from lateral DCN, VAL and cortex through still and move periods. **(L)** Left: LFP dominant frequency for each time bin in (**K**), for still and move periods. Right: Dominant frequency for all time bins of recordings in (**J**) (see **Methods**). All error plotted as +/- SEM. **(M)** Inter-region dominant frequency correlations for time bins in move (top) and still (bottom) conditions. Comparisons shown for DCN vs. thalamus (left) and thalamus vs. cortex (right). Values cluster around the *delta* frequency band during still periods. **(N)** Inter-region *delta* fraction of power for timepoints in move (gray) and still (red) periods. Histograms show probability distributions of single-region *delta* fraction of power.

To interrogate population dynamics in the cerebellothalamocortical system, we simultaneously recorded *in vivo* from the lateral DCN (dentate nucleus, DN), motor thalamus and sensorimotor cortex through movement and still periods (**Figure 1I**). The local field potentials (LFPs) of all three regions exhibited a strong *delta*-band oscillation confined to periods of awake stillness (**Figure 1K**, **1L, Supplemental Figure 1**). We collected the dominant frequencies for each time bin and examined whether these frequencies were shared across these three regions. We found a concordant increase in the *delta* band during stillness and not during movement (**Figure 1M**). Similarly, the *delta* oscillation amplitude shows corresponding increases in *delta* power between regions during stillness (**Figure 1N**). These data indicate that fluctuations in the *delta* rhythm are often shared along the cerebellothalamocortical axis, and *delta* power significantly increases during periods of stillness.

### Cerebellothalamic input is coherent with thalamic *delta* population oscillations

While it is well-established that the LFP correlates with the membrane potential in the thalamus and cortex^20–22^, it is less understood in the DCN, which contain heterogeneous cell populations^23,24^ and project to a multitude of diverse targets^24,25^. To test whether the *delta*-frequency LFP oscillation reflects the aggregate output of DCN neurons onto motor thalamic territories, we performed simultaneous calcium photometry of DCN axons and LFP recordings of the motor thalamus using an optical fiber fastened to a multielectrode array. This “photometrode” approach enabled localized recordings of both postsynaptic and presynaptic populations (**Figure 2A**, see **Methods**). We showed that the activity of DCN axons also oscillates with a *delta* frequency (**Figure 2B, Supplemental Figure 2A-B**). We then compared the timing of each signal’s *delta*-band components. Peak-triggered averages (PTAs) to peaks in thalamic LFP *delta* reveal prominent *delta*-oscillatory activity in DCN axon fluorescence, selectively in stillness (**Figure 2C**, top). Given the limited signal-to-noise of the calcium signal, we confirmed these findings by computing the reverse comparison: the thalamic LFP also shows heightened *delta*-oscillatory activity around DCN peaks in stillness (**Figure 2C**, bottom). Individual experiments had slightly different preferred dominant frequencies within the *delta* band, so we used this variability to test whether the preferred dominant *delta-*band frequency was matched on a per-experiment basis. We found that the DCN and thalamic signals have highly correlated preferred dominant frequency in individual experiments (**Figure 2E**) and that DCN peaks preceded the maximal thalamic LFP PTAs (**Figure 2D**), suggesting that the DCN oscillation is temporally positioned to modulate the thalamocortical oscillation. Comparisons with the calcium-independent, isosbestic control channel and shuffled *delta* peak times did not show similar changes (**Figure 2F, Supplemental Figure 2C**). Consequently, not only do DCN inputs oscillate at *delta* frequencies, but they are phase- and frequency-locked to thalamic *delta* activity. This suggests an important role for the *delta*-band oscillation in cerebellothalamic synaptic transmission during stillness.

**Figure 2:**
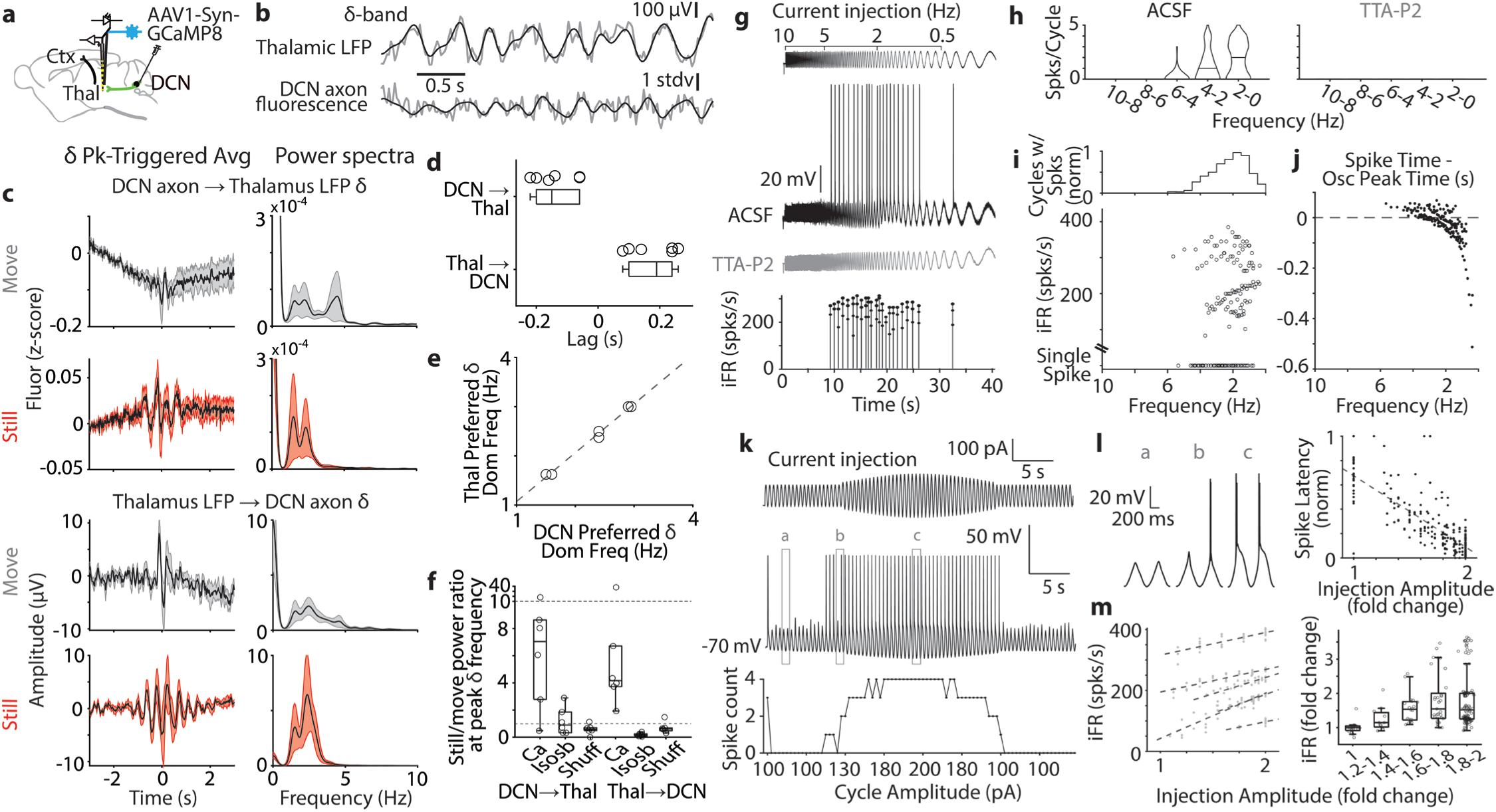
*Delta*-band correlation between cerebellar axon activity and thalamic LFP matches the thalamic frequency preference. **(A-F)** Calcium photometry of cerebellar axons was simultaneously recorded with the thalamic LFP. **(B)** Raw LFP and DCN fluorescence (gray) overlaid with the *delta*-band component (black). **(C)** *Delta*-filtered DCN fluorescence was triggered off the peaks of thalamic LFP (top) and vice versa (bottom). **(D)** Time lags between DCN and thalamic signals generated from (**C**). **(E)** Strong correlation (R-squared: 0.996, p = 2.558e-05) between preferred DCN and thalamic dominant frequencies by insertion (n = 6 insertions in N = 4 animals). **(F)** Ratio of *delta* power in still/move conditions, including isosbestic and shuffled controls. **(G-L)** The frequency preference of thalamic neurons was tested using whole-cell recordings in slice preparation. **(G)** Top: “Chirp” current (±75 pA, sweeping from 10-0.1 Hz) injected into thalamic neurons (n = 6). Middle: Response to current injection in ACSF (black) and CaV3.1 blocker TTA-P2 (grey). Note absence of spikes with blocker in bath. Bottom: Instantaneous firing rate (iFR) of neuron in ACSF. CaV3.1-mediated burst firing occurs between 1-5 Hz. **(H)** Number of spikes per chirp current cycle in ACSF (left) and TTA-P2 (right). Data was binned by cycle frequency. Line denotes median. **(I)** Per-cycle iFR at each chirp frequency. **(J)** Spike times relative to oscillation peak. Spikes occur just before oscillation peak within *delta* band. **(K)** Top: A 2-Hz oscillatory current of varying amplitude was injected into thalamic neurons (n = 6). Middle: Response from example neuron. Bottom: Spikes per burst varies with current injection amplitude. **(L)** Left: Membrane potential in (**K**) showing bursts riding on LTS potential. Right: Normalized latency to spike from previous trough of current oscillation shows linear relationship with injection amplitude (R-sq = 0.78). **(M)** Left: Per-cycle iFR increases with injection amplitude. Lines represent individual cells. Right: Per-cycle iFR fold change from baseline binned by injection amplitude.

### CaV3.1 enables burst encoding of oscillatory inputs

*Delta*-oscillatory cerebellar input might enhance still-period cerebellothalamocortical communication via thalamic resonance in burst mode. To characterize these properties, we performed *in vitro* slice electrophysiology in motor thalamic cells. Current-clamped cells held at hyperpolarized burst-mode potentials were injected with a “chirp” oscillatory current which logarithmically decreased from 10 to 0.1 Hz (**Figure 2G**, top). CaV3.1-dependent burst firing selectively occurred during oscillation cycles within *delta*-band frequencies (**Figure 2G**, bottom). The number of spikes observed per cycle increases within *delta*-band frequencies (**Figure 2H**), as does the initial instantaneous firing rate (iFR; **Figure 2I**). Burst responses within the *delta* band fire preferentially on the rising phase of the input oscillation (**Figure 2J, Supplemental Figure 2D**). These experiments corroborate previous work describing thalamic resonance^6,7^, confirm a frequency preference of thalamic burst firing for *delta*-oscillatory inputs, and suggest that such inputs from DCN can elicit maximal thalamic responses during burst mode.

While it has been established that thalamic neurons have a frequency preference, the capacity for bursts to represent the amplitude of input oscillations is less clear. We injected *delta*-frequency, amplitude-modulated current into thalamic neurons in burst mode, and observed bursts with more spikes (**Figure 2K-L**) at higher rates with increased cycle amplitude (**Figure 2M**). Spike latency shortened with increasing amplitude, corresponding with a larger and shorter-latency LTS (**Figure 2L**). Overall, thalamic burst firing can encode amplitude of oscillatory inputs in a frequency-dependent manner, demonstrating how the long-range *delta* oscillation might act as a carrier frequency for cerebello-cortical communication.

### Burst-mode reliably gates high-frequency synaptic activity

To examine the impact of more realistic cerebellar inputs, we needed to simultaneously stimulate specific DCN inputs in an oscillation frequency- and phase-specific manner while measuring thalamic response. This is not feasible *in vivo*, so we instead performed *in vitro* dynamic clamp experiments in motor thalamic neurons (**Figure 3**). To correctly implement this approach, we quantified the size and number of DCN inputs, the AMPA/NMDA ratio, and short-term synaptic plasticity (**Supplemental Figure 3**, see **Methods**). Our estimates for these parameters are generally in agreement with expectations in the field^26–29^: we interpreted the 5-6 inputs measured in slice as a lower bound for physiological input number and used 10 inputs as an average number of DCN inputs per thalamic neuron, with an average input size of 186 pA (2.86 nS). The DCN single-unit activity from quiescent still mice ^13^ was combined with these properties to construct excitatory conductance streams that simulated DCN synaptic activity (**Figure 3C**, top and middle, see **Methods**) for dynamic clamp. We specifically evaluated how well the dynamic clamp emulated *in vivo* whole cell measurements and previous work^15,26^ (**Supplemental Figure 4A, B**). Membrane variance between 10 and 15 inputs most resembles *in vivo* whole cell variance (**Figure 1H),** corroborating our estimated input numbers and sizes. The inflection of increased spike-triggered postsynaptic potential sizes and spike precision indicated a functional membrane potential transition point for burst/tonic modes (-70 to -75 mV, **Supplemental Figure 4C, D, E**) and matched the transition point observed *in vivo* (**Figure 1E-F).** Together, these experiments indicated that we successfully emulated the major relevant features of the cerebellothalamic synapse.

**Figure 3:**
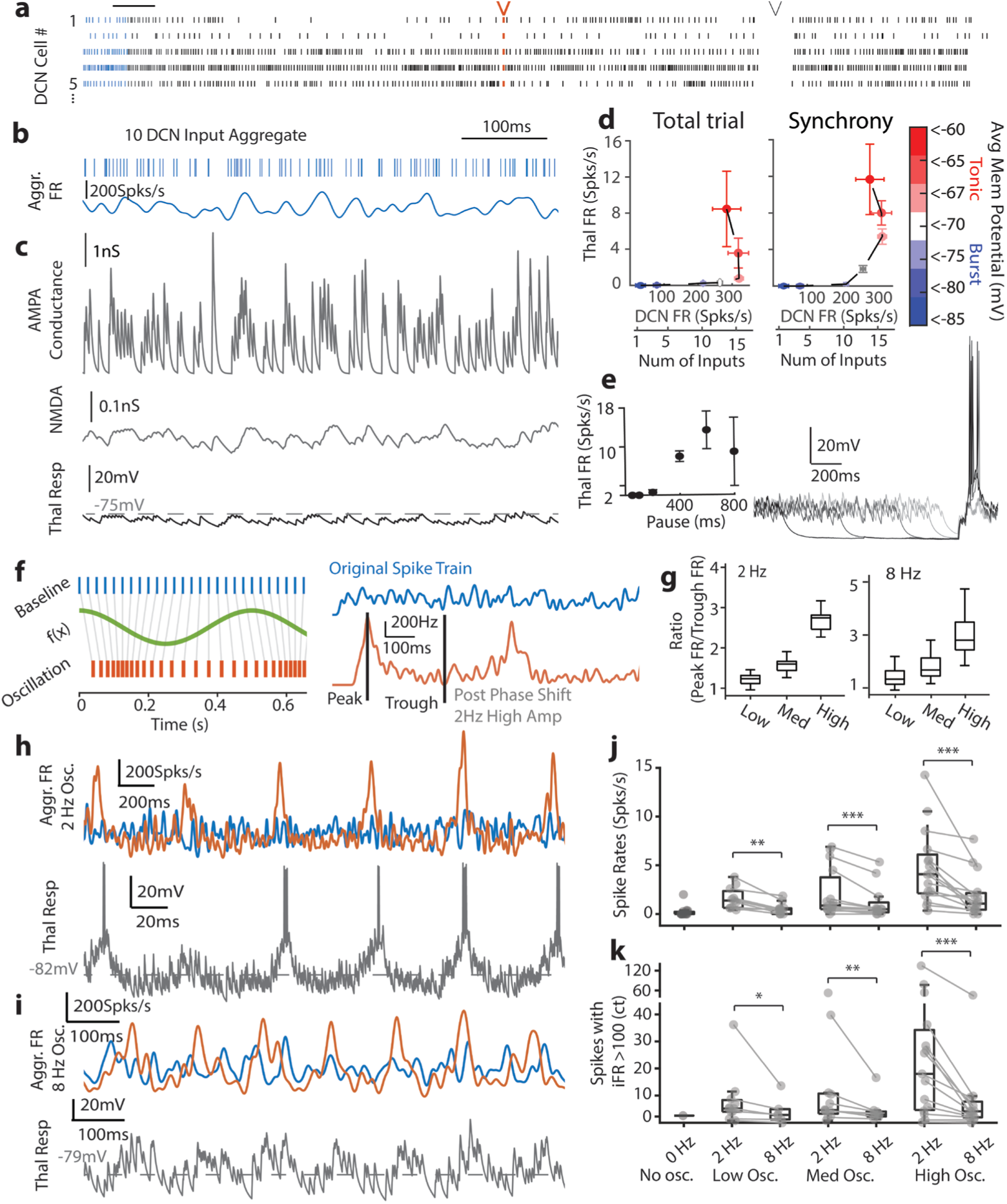
Burst mode gates synchronous spikes and short pauses but preferentially amplifies *delta*-oscillating DCN activity. **(A)** Spike trains from “still” DCN neurons were taken from 10 randomly selected cells^13^ and used in dynamic clamp. A synchronized spike (red arrow) and a pause (black arrow) were inserted into the spike trains to test the role of synchrony and pause on thalamic activity. **(B)** Aggregate activity for the 10 DCN neurons in the first second of (**A**). **(C)** Top: AMPA conductances were calculated from aggregate firing in (**B**). Middle: NMDA conductances for the same ten inputs. Bottom: representative membrane potential response of thalamic neuron when 10 DCN inputs are injected. These conditions rarely resulted in thalamic spiking. Dotted line: average membrane potential of the cell during trial. **(D)** Thalamic firing as a function of input firing rate for the entire trial (left) or for the 100 ms windows when all inputs were synchronized (right, see also (**A**)). **(E)** Left: Thalamic output as a function of pause duration. Right: example traces. **(F)** Left: Phase-shift transformation was performed on DCN single unit activity to simulate an oscillation while maintaining similar overall firing rate from source data (see **Methods**). Right: Resulting firing rates. **(G)** Left: Quantification of the peak and trough ratio after phase shift transformation for 2 Hz oscillation. Right: for 8 Hz modulation. **(H)** Top: aggregate firing rate as in (**B**) (blue) overlaid with 2-Hz oscillatory DCN input firing rate (orange) after phase-shift transformation. Bottom: thalamic response to the 2 Hz oscillatory input. **(I)** As in (**H**) (same cell), but for 8-Hz oscillatory input. Example traces in (**H**) and (**I**) are in response to high oscillation amplitude. **(J)** Quantification of spike rates in response to varying frequency and oscillation amplitudes (n = 18. A subset of cells with matched conditions were used for statistics, see **Table 1**. Paired Wilcoxon rank tests show significant differences between 2 Hz and 8 Hz experiments. Low amp, p = 0.0029. Med amp, p = 4.88e-04. High amp, p = 3.05e-05). **(K)** Number of spikes with iFR > 100 spks/s. (n = 18; paired Wilcoxon rank sum, see **Table 1**. Low amp, p = 0.016. Med amp, p = 0.0078. High amp, p = 4.88e-04)

Since the most commonly ascribed function of thalamic burst mode is to gate inputs^8^, we tested the gating of high-frequency synaptic activity from the cerebellum. We varied the number of simulated DCN inputs so that the aggregated input firing rate ranged from 10 to 350 spikes per second (**Figure 3A, B**), resulting from the summation of a physiological range of input numbers (5-15) and nonphysiological (1 and 20). Varying inputs from physiological to nonphysiological regime revealed the bimodal input transformation. At physiological regimes, motor thalamic neurons were mostly in burst mode (< -75 mV), and few spikes were observed (**Figure 3C**, bottom, **Figure 3D**, left), indicating effective gating maintained by burst mode. When the number of simulated inputs increased to high nonphysiological levels, the membrane potential of the cell entered tonic mode and thalamic firing increased (**Figure 3D**, left). This confirms the capacity of burst mode to gate even high-frequency cerebellar inputs, but contrasts with the substantial, sustained burst-mode activity observed *in vivo* (**Figure 1**).

### *Delta*-oscillating DCN activity can be preferentially amplified by burst mode

To gain insight into the lack of sustained bursting activity *in vitro*, we challenged the gating of burst mode with various patterns of activity in the hopes of replicating *in vivo* observations. A synchronized spike across all inputs in the DCN spike train caused a slight leftward shift in the input-output relationship (**Figure 3D**, right) but did not induce reliable bursts. However, a synchronized pause in conductances longer than 400 ms reliably induced bursts (**Figure 3E**). We reasoned 400 ms pauses might be similar to the troughs of an input oscillating at *delta*-band frequency, which may enable the more sustained bursting activity as seen *in vivo*.

To test the hypothesis that *delta*-oscillating activity can be preferentially encoded and amplified in burst mode, we constructed phase-shifted DCN spike trains (10 inputs) from *in vivo* source data to simulate effects of 2 Hz population oscillation with three different oscillation amplitudes (**Figure 3F, H**). We compared these with a population oscillation of 8 Hz, outside of *delta*, as a control (**Figure 3I**). The firing rates of these population oscillations ranged from ∼1.2x (low) to ∼2.7x (high) in amplitude (**Figure 3G**), which is within the physiological range of DCN dynamics^12–14,30^. When these oscillating inputs were applied, 2 Hz population oscillations induced bursting more frequently than at 8 Hz (**Figure 3H-K**). Consistent with this result, the oscillation cycle-averaged membrane potential showed a boost in burst mode with the 2 Hz but not 8 Hz oscillation (**Supplemental Figure 5A-B**). This evidence indicates that *delta-*oscillating inputs can be represented by thalamic burst mode.

### The cerebellum must phase-lock to thalamic *delta* to enhance thalamocortical transmission

These results suggest that cerebellar inputs might be sufficient to generate a thalamic oscillation. However, it is likely that cortical and reticular feedback circuits can also drive thalamic oscillations^31,32^. To test how cerebellar inputs might constructively or destructively interfere with an ongoing thalamic oscillator, we generated oscillatory activity as in **Figure 3** (**Figure 4A**) but also approximated a thalamic oscillation with a ±25 pA current injected in-phase or antiphase with the 2 Hz cerebellar oscillation (**F**i**gure 4B**). In-phase injections only modestly increased spiking, burst activity (**Figure 4C**) and *delta* power in the membrane potential **(Figure 4D-G**), perhaps owing to saturation of the thalamic output at the higher magnitudes. When current was injected antiphase to the cerebellar oscillation, spiking, bursting, and *delta* power reliably decreased (**Figure 4C-G**). This indicates that proper phase-locking to any ongoing thalamic oscillation is essential for cerebellothalamocortical transmission.

**Figure 4.**
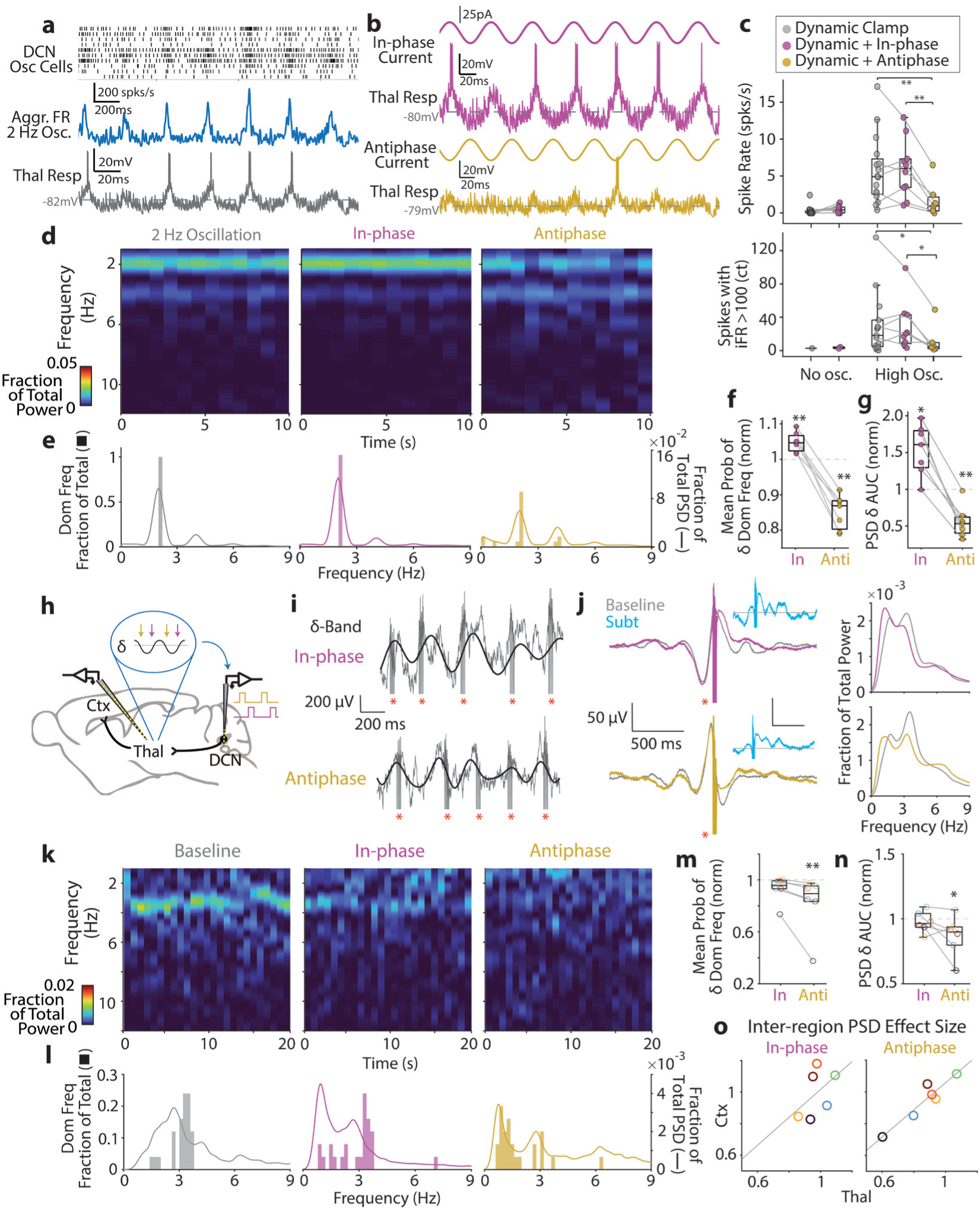
Phase-matching cerebellar and thalamic oscillations enables cerebellocortical transmission. **(A)** Top: Oscillating DCN spike train with 10 inputs for dynamic clamp. Middle: Aggregated firing rate for above. Bottom: Example thalamic response to the oscillating input. **(B)** Top: 2 Hz oscillating current injected in-phase with the oscillating DCN spike train. Bottom: 2 Hz oscillating current injected antiphase with the oscillating DCN spike train. **(C)** Quantification of spike rates (top) and burst counts (bottom) in response to current injection (paired Wilcoxon rank sum, see **Table 1**. Spike rates: 2 Hz vs antiphase p = 0.0078; in-phase vs antiphase p = 0.0078. Spikes with iFR>100: 2 Hz vs antiphase p = 0.031; in-phase vs. antiphase p = 0.015). **(D)** Example spectrogram of thalamic membrane potential with cerebellar oscillatory conductances (left), with in-phase current (middle), and with antiphase current (right). **(E)** Bars: Dominant frequency histogram of recording durations in (**D**). Lines: averaged PSD across all cells (n = 7). **(F)** Mean probability of dominant frequency histograms within *delta* band, for in-phase and antiphase conditions in all recordings. Values are normalized to 2 Hz oscillation trials (paired Wilcoxon rank sum, in-phase p = 0.0078; antiphase, p = 0.0078). **(G)** Normalized area-under-curve (AUC) values for averaged PSD across all cells (paired Wilcoxon rank sum, in-phase p = 0.016; antiphase, p = 0.0078). **(H)** Triple-site recordings were performed as in Figure 1 (n = 5 insertions in N = 3 animals). Electrodes stimulated the DCN on the rising phase (in-phase, magenta) or falling phase (antiphase, gold) of the *delta*-filtered thalamic LFP. **(I)** Alignment of stimuli on raw (gray) and *delta*-band component (black) of thalamic LFP. Red asterisks denote stimulus artifacts. **(J)** Left: STAs of thalamic LFP for in-phase and antiphase conditions. Baseline STAs were generated from similar timepoints extracted from no-stimulus recordings (see **Supplemental** Figure 6A-C**, Methods**). Inset: baseline-subtracted STAs (in-phase max amplitude 37.1 μV, antiphase max amplitude 28.7 μV). Right: PSD plots of STA traces. **(K)** Example spectrograms during baseline, in-phase and antiphase stimulation recordings. **(L)** Bars: Dominant frequency histogram of recording durations in **(K)**. Lines: Averaged PSD across all recordings (n = 6). **(M)** Mean probability of dominant frequency histograms within *delta* band, for in-phase and antiphase conditions in all recordings (unpaired Wilcoxon rank sum: in-phase vs.1 p = 0.9935; antiphase vs. 1 p = 0.0011). Values are normalized to baseline. **(N)** Normalized AUC values for thalamic LFP in in-phase and antiphase conditions (unpaired Wilcoxon rank sum: in-phase vs. 1 p = 0.8680; antiphase vs. 1 p = 0.0238). **(O)** Normalized PSD AUC values for thalamic vs. cortical LFP. Lines depict linear fit (in-phase R = 0.23, antiphase R = 0.89).

We next tested whether the pattern of cerebellar activity could modulate thalamic *delta in vivo*. If phase-locking to thalamic *delta* is necessary for cerebellothalamic transmission, we hypothesized that inducing cerebellar activity antiphase to thalamic *delta* would be sufficient to diminish the thalamocortical *delta* rhythm. Conventional loss/gain-of-function approaches (i.e. using optogenetics/chemogenetics) would test the role of broad changes in cerebellar activity but lack the temporal specificity to investigate phase-dependence. Therefore, we performed closed-loop experiments in which the *delta* component of the thalamic LFP during still periods was used to trigger DCN electrical microstimulation (**Figure 4H**). Electrodes were arranged to record from the DCN, motor thalamus and sensorimotor cortex as in **Figure 1**, and DCN stimuli (5-10 µA) were set to be delivered either on the rising (in-phase) or falling (antiphase) portions of the thalamic LFP (**Figure 4H-I, Supplemental Figure 6A-C**). Consistent with dynamic-clamp experiments (**Supplemental Figure 5A**), the thalamic response depended on the phase of the field potential (**Figure 4J**): rising, in-phase stimulation resulted in slightly larger field responses (10 µV) and different frequency components than the antiphase stimulation. Throughout the course of the stimulation period, both in-phase and antiphase stimulation resulted in more complex, leftward-shifted power spectra (**Figure 4J**, right). In-phase stimulation minimally altered total *delta* power, but antiphase stimulation was effective at diminishing *delta* (**Figure 4K-N**), in line with dynamic clamp experiments (**Figure 4F-G**).

We wondered whether this stimulation would be sufficient to modulate cortical *delta*. We found a strong linear correlation between the antiphase effects on the thalamus and the cortex (**Figure 4O**), indicating that inter-region communication is also amplified by cerebellar phase-matching with the thalamic *delta* carrier frequency. The relationship between in-phase effects on the thalamus and cortex was weaker, likely owing to the overall weak effects on the thalamic *delta* with in-phase stimulation. These results indicate that cerebellar phase-locking to the thalamic *delta* enables maximal cerebellocortical transmission during still periods.

### Thalamocortical *delta* rhythms are dependent on CaV3.1 in thalamic neurons receiving cerebellar input

We next evaluated the role of the *delta* carrier frequency in cerebellothalamocortical transmission using loss-of-function experiments. To do so, we designed an AAV viral construct containing a Cre-dependent CRISPR-Cas9 scaffold targeted to *Cacna1g,* the gene encoding CaV3.1 (**Supplemental Figure 7A-B**); transfection into Neuro2a cells successfully mutated *Cacna1g* in the presence of Cre and SaCas9/*Cacna1g* targeting gRNA (**Supplemental Figure 7C**). We then injected the CRISPR-Cas9 virus into the motor thalamus and an AAV1-Cre virus into the DCN, using the anterograde properties of the AAV1 serotype at high titer^33^ to selectively impair *Cacna1g* function in cerebellar-recipient motor thalamic neurons (**Figure 5A, Supplemental Figure 8A**). *In vitro* electrophysiological assessment revealed a significant proportion of cells with an impaired LTS and/or burst phenotype (**Supplemental Figure 7D-E**).

**Figure 5:**
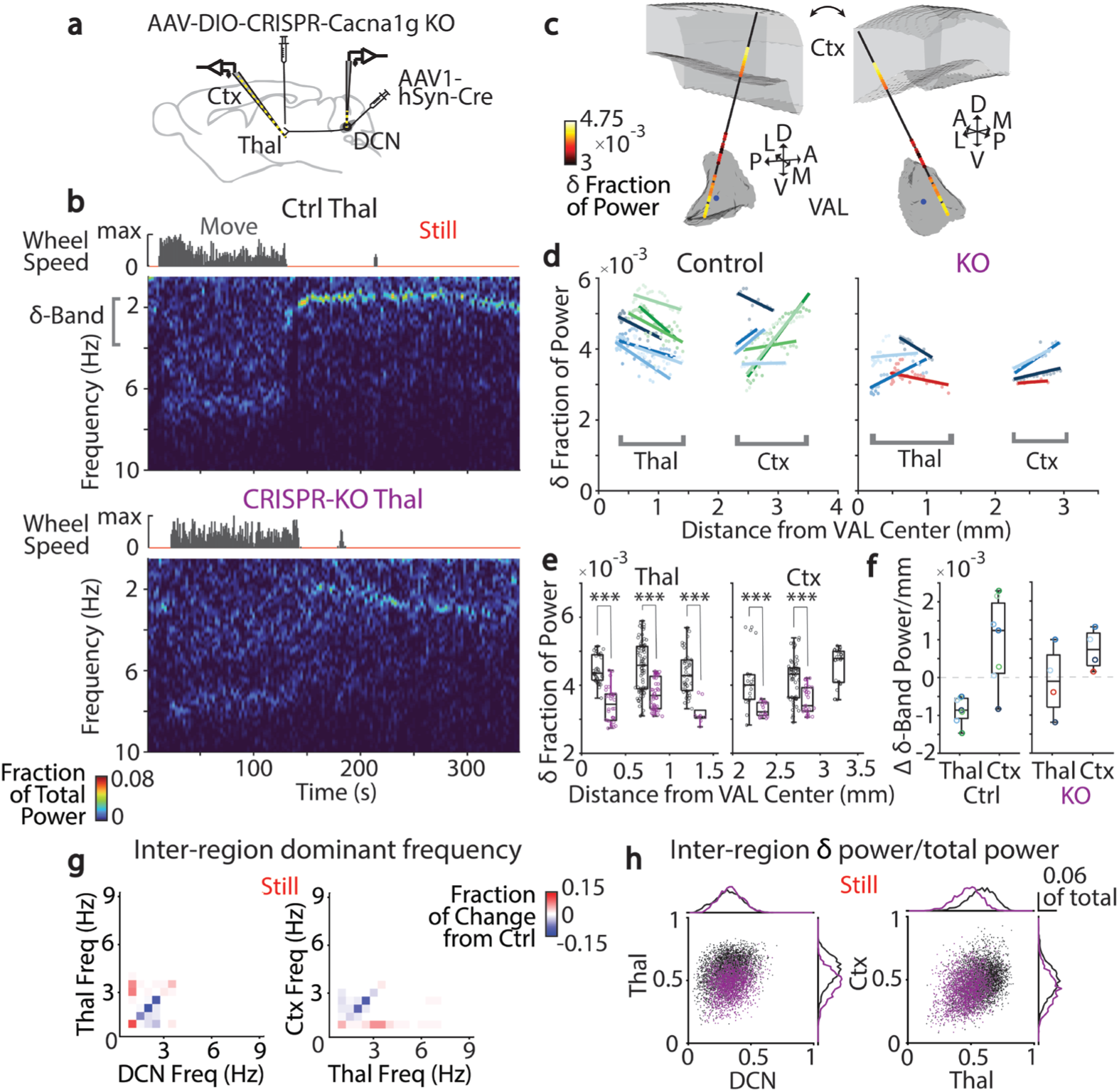
Viral CaV3.1 knockout in VAL neurons receiving cerebellar input impairs *delta*-band oscillations. **(A)** CRISPR-Cas9 was used to knock out CaV3.1 selectively in DCN-receiving thalamic cells (see **Methods**). Triple-site recordings were performed as in Figure 1. **(B)** Top: Spectrogram of thalamic LFP in noninjected (control) animal. Bottom: similar recording from KO animal. Examples were channel impedance- and location-matched (see **Supplemental** Figure 7F). **(C)** Reconstruction of sample probe insertion. Spheres depict locations and still-period *delta*-band power of probe channels. **(D)** Average still-period *delta*-band power for all channels (dots) and linear regression fits per insertion (lines) in thalamus and cortex. Distance to VAL weighted centroid was computed from a normalized brain. Color families distinguish recordings made with same probe; hues within each family represent individual insertions (control, n = 7 insertions, N = 3 animals; KO, n = 4 insertions, N = 2 animals). **(E)** Still-period *delta*-band power binned by distance from VAL. Gray: control channels; violet: KO channels. Asterisks indicate statistical significance (paired Wilcoxon rank sum, see **Table 1**). **(F)** Slopes of linear fits in (**D**) for control and KO recordings. **(G)** Inter-region correlations of dominant frequency as in Figure 1M; colors represent fraction of change of KO from control. **(H)** Inter-region *delta* fraction of power for still-period timepoints in control (black) and KO (violet) animals.

We hypothesized that impairing CaV3.1-mediated potentials would diminish *delta*-oscillatory activity in the thalamus, so we performed *in vivo* recordings of DCN, thalamic and cortical LFPs in mice with CaV3.1 knocked out (KO) in thalamic neurons receiving cerebellar inputs (**Figure 5A**). KO animals displayed a marked decrease in the still-period thalamic *delta*-band oscillation (**Figure 5B**). To quantify this change, we calculated the average *delta* power for each thalamocortical electrode channel in stillness (**Figure 5C**). In control animals, *delta* power increases with proximity to center of motor thalamus (**Figure 5D**, left; **5F**, left). Similarly, *delta* power in cortical channels increased with distance from the lateral ventricle (**Figure 5E**, right), consistent with the distribution of thalamocortical terminals predominantly in cortical layer V and corticothalamic pyramidal neurons in layers V and VI^34–36^. In contrast, KOs demonstrated weaker baseline thalamocortical *delta* power (**Figure 5E, Supplemental Figure 7G-H**) and reduced location-dependent changes (**Figure 5F**). Remarkably, *delta*-band frequency and amplitude inter-region correlations were also diminished in KO animals across the thalamus *and* cortex, highlighting the importance of these neurons for generating the cortical *delta* rhythm (**Figure 5G**, **5H**). Importantly, KO DCN *delta* power resembled control (**Figure 5H, Supplemental Figure 7G,** left), indicating that while thalamocortical *delta* is disrupted, cerebellar outputs remain largely intact. In summary, thalamic neurons receiving cerebellar input are critical mediators of *delta*-band thalamic and cortical oscillations.

### Selective CaV3.1 knockout in cerebellar thalamus impairs motor learning

The content and significance of information transmitted by the *delta* carrier frequency remains unknown, so we investigated the impact of *delta*-signal dysfunction on motor behavior. We reasoned that if *delta* rhythms were more typically observed during periods of stillness, impairing *delta* should minimally affect “online” functions like motor coordination. In agreement with this, no overt motor deficits were observed in KO animals (**Supplemental Video 1+2**). To quantify this, we investigated the baseline motor behavior of KO animals and age-matched controls (sham-injected and control vector-injected, see **Methods**) using the Blackbox gait analysis system, which quantifies the kinematics of each limb during spontaneous bouts of movement. Gait metrics showed minimal differences between controls and KOs (**Figure 6A-C**, **Supplemental Figure 8A-D**).

**Figure 6.**
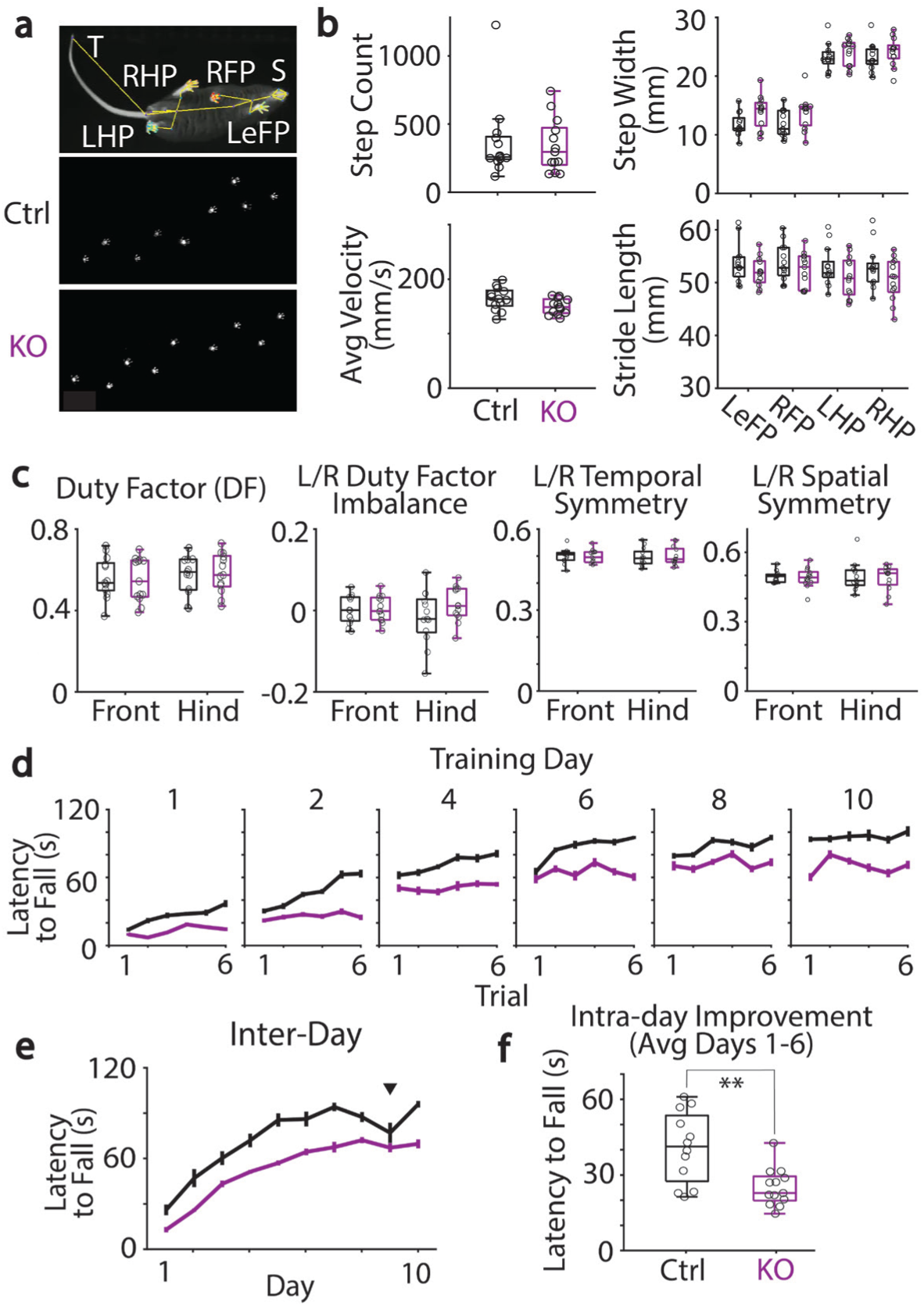
Viral CaV3.1 knockout in VAL neurons receiving cerebellar input produces motor learning deficits but preserves baseline motility. **(A)** Blackbox software-detected body-position skeleton (top) and frustrated total internal reflection (FTIR) measurement of sample movement bout (middle, bottom). LeFP = left front paw, RFP = right front, LHP = left hind, RHP = right hind, S = snout, T = tail. **(B)** All statistical analysis for gait parameters used the FDR-corrected Kruskal-Wallis test. Top left: Total step count for control (black; sham-injected N = 6, empty vector-injected N = 7) and KO (violet, N = 13) animals, used to define movement bouts (KO bout range 10-42, mean = 17.625, control bout range 9-27, mean = 17.333; p = 0.9795). Bottom left: Average walking velocity per bout (p = 0.4121). Top right: Average step width by paw (LeFP p = 0.4121, LHP p = 0.5712, RFP p = 0.4121, RHP p = 0.5712). Bottom right: Average stride length by paw (LeFP p = 0.5712, LHP p = 0.5712, RFP p = 0.5712, RHP p = 0.5712). **(C)** Left to right: front and hind paw duty factors as fraction of time spent on floor (front p = 0.9795, hind p = 0.9795). Duty factor imbalance between contralateral paws (front p = 0.1967, hind p = 0.9795). Temporal symmetry between contralateral paws, measuring relative swing phase time (front p = 0.9795, hind p = 0.5712). Spatial symmetry between steps taken with contralateral paws, measuring relative step placement (front p = 0.9795, hind p = 0.9795). **(D)** Intra-day mean latency to fall across training days of accelerating rotarod task (3-way ANOVA, trial (first vs. final) interaction with experimental group, p = 0.0058, see **Table 1**). **(E)** Inter-day mean latency to fall on rotarod task (2-way ANOVA, control vs. KO p = 0, see **Table 1**). Arrow denotes day of intermittent construction noise below the behavior room, disturbing experiments. **(F)** Average intra-day performance improvement (max latency to fall-first trial) during task acquisition period on days 1-6 (unpaired Wilcoxon rank sum, control vs. KO p = 0.0039).

Instead, we might expect dysfunction in motor computations that occur “offline,” such as consolidation of learning. So, we next tested control and KO mice on a motor learning task that requires cerebellothalamocortical processing: the accelerating rotarod^37–40^. Mice were tested in 6 rotarod trials per day for 10 days, with an intertrial interval of 2 minutes (see **Methods**). KOs displayed reduced task performance across all training days (**Figure 6D-E**). Closer scrutiny of their motor performance indicated that KOs displayed minimal intra-day improvement (**Figure 6D, F**, **Supplemental Figure 8E**). This indicates that *delta* during stillness is essential for trial-to-trial motor learning, and might function as a post-trial consolidation mechanism. KOs did exhibit improvement between successive training days (**Figure 6E**), pointing to preserved mechanisms of learning at longer intertrial intervals. In summary, we found that KO of CaV3.1 in thalamic neurons receiving cerebellar input produced specific learning deficits, most prominently on a short-term basis, while largely preserving baseline motor behaviors. These results suggest a role for the cerebellothalamocortical *delta* oscillation in transmitting information crucial for optimal motor learning.

## DISCUSSION

Here we identify a coherent *delta* oscillation in activity across the cerebellothalamocortical axis when mice are still. We show that the mechanism of transmission across this axis resembles a carrier frequency, as it facilitates transmis ‘sion across long distances (**Figure 1)**, represents input frequency and amplitude (**Figure 2-3**), and amplifies or suppresses inputs based on frequency and phase (**Figure 3**–**4**). It is dependent on CaV3.1, impacts cortical activity, and is critical for motor learning (**Figure 5**–**6)**.

We show that cerebellar inputs tune into a thalamic *delta* oscillator and can constructively or destructively modulate *delta* via timing of these inputs (**Figure 3**–**4)**. We suspect that the convergence of basal ganglia inputs^29,37,41^, cortical inputs^27,42^, and reticular thalamic inputs^43–45^ at the thalamic territories receiving cerebellar input can also support or interfere with the thalamic *delta* oscillator. The timing of these inputs is key to how faithfully they are transmitted out of the thalamus. We do not know what signals appropriately time the cerebellar *delta* oscillator, though CaV3.1’s presence across the olivocerebellar loop^46–49^ and the many routes by which mossy fibers might relay cortical timing to the cerebellum^50,51^ are promising candidates.

We provide evidence that this *delta* carrier frequency is important for motor consolidation, which aligns strongly with recent work examining offline motor learning and replay^38,39^. This work highlights the importance of cerebellar computations in offline motor learning. They posit that reactivation of locomotor-related neural activity during quiescent intertrial intervals reflects the exploration of strategies critical for appropriate motor learning. Our observations indicate that *delta* might be essential to transmit that neural activity throughout the thalamocortical axis during those quiescent periods.

Extensive work in cortical and hippocampal physiology has generated frameworks for oscillation-based communication mediated by population coherence and resonance^1–5^. These works have characterized transient interregional coherence of *gamma* and *theta* oscillations to facilitate short-range communication in sensory pathways. In complement to this, *delta*-dependent cerebellothalamocortical transmission is best described as a “carrier frequency”: it utilizes *delta* as a continuous long-range carrier to amplify relevant information, exploits intrinsic thalamic subthreshold oscillations and resonance, and uses thalamic CaV3.1’s impressive nonlinearity. LTS events range from single spikes to quintuplets and beyond, and are ideal for distributing well-timed information broadly across the cortex^34^. Finally, the carrier frequency framework suggests that several streams of afferent information can be simultaneously encoded and deliberately “tuned” to by a receiver, as might be the case during changes in thalamic firing mode^42^.

Beyond motor circuits, the high density of CaV3.1 across both first-order and higher-order thalamus^47,52^ suggests that *delta* might play a role in many transthalamic pathways, particularly in quiescent conditions when basal activity is low and boosts in transmission are needed. Indeed, cerebellothalamic projections extend beyond the motor regions, specifically to higher-order thalamus^24,53^. It is also worth noting that high levels of single-cell activity are not necessary for this mechanism to operate—high input convergence may also assemble into oscillatory conductances, as might be seen in cortical inputs to the higher-order thalamus, or inhibitory basal ganglia inputs. The *delta* mechanism and its role in learning might thus extend to many sensorimotor modalities.

## METHODS

### Animals

All animal procedures were carried out in accordance with the NIH and the Institutional Animal Care and Use Committee (IACUC) guidelines and protocols approved by the Pennsylvania State University College of Medicine IACUC (protocol #PROTO202202365) or by the Marine Biological Laboratory Institutional Animal Care and Use Committee. C57BL6/J mice of either sex from Jackson Laboratory were used for all experiments.

### General surgery protocol

Mice were anesthetized and maintained under 2% isoflurane; body temperature was maintained throughout surgery with a heating pad. Mice were secured to a stereotaxic surgery instrument (Stoelting). Eye lubricant was applied and reapplied throughout surgery as needed. The skin of the head was depilated and sanitized, and an incision was made to expose the cranium. The anterior-posterior (AP) and medio-lateral (ML) axes of the cranium were leveled using bregma and lambda suture points and points equidistant from bregma, respectively. At the end of surgical procedures, mice were subcutaneously administered slow-release analgesic (0.1 mL Ethiqa-XR, Midwest Veterinary) and monitored for the following 5 days.

### Viral injections

A craniotomy was made at the desired coordinates. A glass pipette containing the viral solution and attached to an injector (Nanoliter 2020, WPI) was slowly lowered into the brain to the desired dorso-ventral (DV) location. Viruses were injected at a rate of <3 nL/s. The micropipette remained in place for at least 5 min following injection before being slowly retracted from the brain tissue. Incisions were closed with sutures. Viral expression was allowed to proceed for 2-3 weeks before experimentation. Recombinant AAV (rAAV) vector genomes containing AAV2 ITRs were packaged using the indicated capsids; vectors are subsequently referred to by their capsid identity (e.g., rAAV9 or rAAV-DJ/8). For acute slice synaptic property and initial *in vivo* LFP characterization, mice at 3 weeks of age were injected with 400 nL of rAAV9-hSyn-hChR2(H134R)-EYFP (Addgene 26973, titer 1.9 x 10^13^ gc/mL or for *in vivo* experiments, UPenn Vector Core rAAV1-Syn-ChR2(H134R)-YFP) in the bilateral deep cerebellar nuclei (AP: -6.0, ML ±2.25, DV: -2.5 mm).

For photometry experiments, mice were injected in the deep cerebellar nuclei (coordinates as previous) with 200-300 nL of either rAAV1-syn-jGCaMP8f-WPRE (Addgene, 162376, titer ∼2 x 10^13^ gc/mL) or rAAV1-syn-jGCaMP8m-WPRE (Addgene, 162375, ∼2 x 10^13^ gc/mL).

For CRISPR-KO experiments, mice were injected at 4-6 weeks old. Viral solutions were injected bilaterally at the interposed nucleus (IP; AP: -6.15, ML: ±1.84, DV: 2.61 mm), dentate nucleus (DN; AP: -5.84, ML: ±2.36, DV: 2.51 mm), and ventral anterior-lateral thalamus (VAL; AP: -1.18, ML: ±1.13, DV: 2.64 mm). Both IP and DN were injected (200 nL/site) with a mixture of rAAV1-hSyn-Cre-WPRE.hGH (Addgene 105553, titer 2.4 x 10^13^ gc/mL) and a 1:100 dilution of rAAV-rh10-Ef1a-EYFP-DIO (3.42 x 10^13^ gc/mL). VAL was injected (300 nL/site) with a mixture of rAAVDJ8-U6 *Cacna1g* gRNA-CMV-SaCas9-DIO (∼ 1.0 x 10^13^ gc/mL) and 1:100 dilute rAAV-rh10-Ef1a-EYFP-DIO (3.42 x 10^13^ gc/mL). Sham injections proceeded as experimental injections, except in VAL, a similar control AAV, rAAVDJ8-U6 gRNA-CMV-SaCas9-DIO virus (∼1.0 x 10^13^ gc/mL) containing an empty gRNA scaffold was injected (see **Viral Construction and Validation**). For sham injections, a pipette containing no viral solution was lowered into the brain at the desired coordinates and then retracted.

### Head-fixed recording implantation

After the initial general surgery protocol, the skull was scored to assist adherence of adhesives. Locations of intended craniotomies were marked on the skull surface. A screw and ground pin were inserted over the left hemispheric cerebellum (AP: -5.5, ML: -2.0). Two additional screws were inserted through the skull (AP: -5.5, ML: ±2.0) to stabilize a steel head bracket. The skull surface, screws, and ground pin were sealed with cyanoacrylate glue. The bracket and scalp were secured to the skull using dental cement (MetaBond). Two points of known DV depth difference were marked in visible locations atop the implant for leveling the skull’s AP axis in later experiments. The craniotomy locations were covered by Kwik-Cast sealant (WPI).

### *In vivo* whole cell recordings

Mice were implanted as for head-fixed recording as above. Mice were minimally acclimated to the recording setup (3 sessions of <1 hour) to limit spontaneous running. On the day of recording, a craniotomy was made above the VAL (AP: -1.2, ML: ±1.00 mm). After at least 3 hours of recovery, mice were head-restrained over a running wheel. Whole-cell recordings were achieved using standard approaches^54^ using 4-7 MΩ pipettes. The Optopatcher (A-M Systems) was used to stimulate cerebellar axons and elicit LFP responses to assist in identifying VAL. There were several important considerations for stable recordings: first, extra pressure and speed were applied when crossing through ventricles and seals were only attempted if mice were not moving. Seal success was greatly improved by maximizing pulsation magnitude before switching to negative pressure. Recording quality was assessed by the presence of overshooting action potentials, and a stable resting potential. Movement on the wheel was recorded using an Arduino reporting the movement of an optical mouse positioned on the side of the wheel.

### *In vivo* multielectrode probe recordings

Optimal probe trajectories based on skull size and intended recording targets were generated using the Neuropixels Trajectory Explorer interface (https://github.com/petersaj/neuropixels_trajectory_explorer, by Andy Peters). For Ctx-Thal recordings, probes were inserted at AP: 0.04, ML -1.25, DV: -4.15, and positioned at 180° azimuth, 19° elevation; for DCN recordings, probes were inserted at AP: -6.3, ML: - 2.5, DV: -2.8, 0° azimuth and 12° elevation. Coordinates were adjusted for cranium size as measured by the distance between bregma and lambda suture points.

Following surgery, mice were acclimated to the recording setup as above. On the day of recording, animals were anesthetized with isoflurane before craniotomies were drilled at the previously marked locations. Animals were administered carprofen for analgesia (0.1 mL) and given at least 3 hours to recover prior to recording.

At the time of recording, animals were head-fixed to a running wheel. Movement bouts were recorded by video cameras (2.0 Megapixel USB Camera, ELP) and a rotary encoder with an Arduino Uno reporting wheel rotation. The cranium was leveled as guided by the marks made during implantation surgery.

*In vivo* electrophysiological data was acquired with an RHS Stim/Recording Controller (Intan Technologies) at a sampling rate of 30 kHz, with frequency bounds set to 0.51 Hz-7.60 kHz. Acute electrophysiological recordings were made using multichannel silicon probes (Ctx-Thal: 64-channel linear L3 probe; DCN: 32-channel linear H3 probe, Cambridge Neurotech). Probes were coated with fluorescent lipophilic dye solutions (Thermofisher, V22889) for post-hoc identification of tract trajectory. Probes and headstages were mounted on and manipulated by a Multi-Probe Micromanipulator System (New Scale Technologies). Probes were lowered near to the craniotomy and inserted into the exposed brain tissue. The initial depth reading at the brain surface was used to determine the ideal insertion depth. Probes were inserted into the brain to the ideal depth at a speed of 3.5 um/s. To ensure movement periods of sufficient length for spectral analysis, the running wheel was manually turned for randomly determined 2-minute durations during the recording to induce movement of the animal. After recordings, probes were retracted from the brain at the same speed. Craniotomies were covered by silicone sealant (KWIK-SIL, WPI) between successive days of recording.

*In vivo* CRISPR-KO recordings were performed on animals that had previously undergone behavioral testing. Animals that exhibited performance deficits representative of the cohort median were selected for electrophysiological testing. Viral expression was verified following conclusion of experiments.

For closed-loop electrical stimulation experiments, the strength of the *delta* oscillation was used to select optimal recording and stimulation channels. A single recording channel in the thalamus was lowpass-filtered using a simple RC circuit with a cutoff frequency of 7 Hz. This signal was used to trigger electrical stimulation from probe contacts in the DCN. Stimulation was set to trigger either on the rising phase (for in-phase experiments) or falling phase of the input signal (for antiphase experiments). A stimulus train of four biphasic 200-µs pulses (cathodic then anodic current, equal amplitude, each 100 µs) at 100 Hz was given at 1-3 selected DCN contacts for a total train duration of 40 ms. A minimum refractory period of 200 ms was imposed between trains to ensure stimulation remained within *delta*-band frequencies. Stimulation amplitude was between 5-10 µA. Control recordings without stimulation were made prior to stimulation experiments to measure baseline *delta* between successive recording sessions.

### Simultaneous electrophysiology and photometry recordings

Multielectrode/optical fiber probes (“photometrodes”) were constructed by cementing (Metabond) an optical fiber (200 µm diameter, 0.37 NA, Neurophotometrics) to a 32-channel linear probe (M2 probe, ML Precision). The optical fiber was angled towards the electrode shaft (∼5°), and the tip was separated from the electrode shank by ∼200 µm and from the electrode tip by ∼1 mm. The size of the optical artifact induced on the electrode contacts in bath ACSF was used to determine which contact was most ideally positioned to simultaneously measure the thalamic LFP and imaging area. This artifact was insignificant once the photometrode was inside neural tissue. This contact was positioned to the requisite recording site in VAL. Lipophilic dye could not be used to coat the electrode to determine recording locations in these experiments because dye fluorescence would be detected by the photometry. Instead, the characteristic rise of fluorescence signal that occurred when entering VAL in a successfully injected animal was used to position the photometrode for recordings.

Electrophysiology data was acquired on an Intan RHD recording system (Intan Technologies) and initially acquired at 30 kHz (0.1–10 kHz bandpass). Photometry data was acquired on the FP3002 (Neurophotometrics) at 50 frames per second for isosbestic and 470 nm channels with an 8-ms integration time per frame. Illumination intensity varied between 0.1-0.5 mW, as measured from the end of the patch cable. Recordings were thus performed using the two-wavelength stimulation (multiplexed mode) in which the isosbestic or 470 nm channel was active every other frame. Fluorescence was recorded using Bonsai^55^. Bleaching was corrected offline by subtracting a 10-minute moving average from the fluorescence signal. Electrophysiology data was resampled to the photometry acquisition rate for all comparisons and analysis. Recordings were conducted in the dark to minimize incidental light.

### *In vitro* electrophysiology

Mice ranging from P35 to P50 (n=64) were anesthetized with isoflurane, then 0.4 mL ketamine/xylazine and transcardially perfused with cold NMDG cutting solution (in ice bath) containing in mM: 92 NMDG, 2.5 KCl, 1.25 NaH_2_PO_4_, 30 NaHCO_3_, 20 HEPES, 25 Glucose, 2 Thiourea, 5 Na-Ascorbate, 5 Na-Pyruvate, 0.5 CaCl_2_-2H_2_O, 10 MgSO_4_-7H_2_O. Solution was titrated with 5M hydrochloric acid until pH level maintained at 7.2-7.3, then oxygenated with 95% O2 / 5% CO2. Coronal slices were made (200 µm) using a vibratome (7000smz-2, Campden Instruments) in cold NMDG solution. Slices were then transferred to a holding chamber with warm ACSF (34°C) containing in mM: 127 NaCl, 2.5 KCl, 1.25 NaH_2_PO4, 25 NaHCO_3_, 25 glucose, 1.5 CaCl_2_, 1 MgCl_2_ and were recovered at 34°C for 30 minutes before being moved to room temperature until recordings began. The osmolarity of cutting and ACSF solution was adjusted to 310-320 mOsm. The flow rate in bath was ∼2 ml/min.

Voltage-clamp recordings were made across the ventrolateral thalamus. Borosilicate glass electrodes (2-4 MΩ) were filled with an internal solution containing in mM: 115 CsGluc, 25 TEA-OH (40%), 10 HEPES, 0.2 EGTA, 5 QX-314-Cl, 4 NaCl, 2 MgATP, 0.4 Na_3_GTP, 10 Na_2_Phosphocreatine. The osmolarity was adjusted to 290-300 mOsm. Thalamic neurons were held at -60 to -70 mV. All experiments were performed at 34-36°C.

In a subset of experiments, 2.5 µM NBQX and 2.5µM CPP were sequentially applied to block AMPRs and NMDARs and isolate input currents. In a subset of experiments, ChR2-expressing DCN axons were stimulated with 473 nm light from an LED in the entire field of view at least 1 mm away from the soma (1ms, 1 mW/mm^2^) for at least 10 trials to obtain a maximal current amplitude. To find the single-input size, the LED light path was restricted to 30% of full field of view, and stimulation proceeded until a location with an “all or none” (∼50%) response was observed.

Due to limitations of ChR2 kinetics, prolonged DCN activity at high frequency (>50 Hz) cannot be measured with optogenetics. ChR2 was used to elicit responses from DCN axons which were subsequently electrically stimulated. With this approach, the optical stimulation of axons occludes electrical stimulation of the same fiber, due to the refractory period of action potentials. In contrast, if the electrically stimulated axon is not expressing ChR2, the optically evoked response does not occlude the electrically evoked response^56^. Optical stimulation was done as in prior experiments. A glass monopolar stimulus electrode (4-5 MΩ) filled with ACSF was placed 2 mm away from the soma to evoke postsynaptic currents. Trials of single optical, electrical or closely timed (1 ms) paired optical/electrical stimuli were conducted until the electrically evoked component could be occluded by optical stimulation. A train of high-frequency electrical stimulation was then delivered. Electrical trains were constructed by concatenating a 30 Hz baseline period, stationary period, and walk period, among which the stationary and walk period were representative spike timing from *in vivo* recordings of DCN neurons when mice were still or moving, respectively^13^. EPSC depression was measured and calculated in an inter-spike-interval (ISI) dependent manner.

### Dynamic clamp

For dynamic clamp experiments, borosilicate glass electrodes (2-4 MΩ) were filled with internal solution, containing in mM: 130 K-Gluconate, 10 Na-Gluconate, 10 HEPES, 10 Phosphocreatine-di(tris) salt, 4 MgATP, 0.3 NaGTP, 4 NaCl. The osmolarity of internal solution was adjusted to 290-300 mOsm. We used the same internal solution as *in vivo* experiments, and corrected for a calculated liquid junction potential of 15 mV *in vivo* and *in vitro*^57^. Cells were held at -75 to -85 mV between trials. All experiments were performed at 34–35°C.

Dynamic clamp was implemented with the dPatch amplifier (Sutter Instruments) which includes integrated capability to perform dynamic clamp^58^. The unitary current waveform was empirically calculated from the mean AMPA/NMDA responses (**Supplemental Figure 3**). The unitary current was then converted to unitary conductance for both AMPA and NMDA-mediated excitation using a reversal potential of -65 mV. We used the two output channels of the dPatch to calculate AMPA and NMDA conductances. For the NMDA conductance stream, a function (1) was applied to adjust the amplitude of the calculated current based on the membrane potential (V)^59^, as part of the dPatch dynamic clamp model.

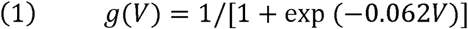

To generate DCN conductance streams, ten-second segments of DCN single unit activity from still mice were used as source data^13^. These inter-spike intervals (ISIs) were used to construct streams of excitatory conductance based on the calculated unitary conductances above. To approximate the results of short-term presynaptic plasticity, the conductances were scaled in an ISI-dependent manner, using the ISI-depression relationship calculated from the electrical stimulation experiments (**Supplemental Figure 3**).

To aggregate DCN population activity, each 10 s conductance was used as an individual DCN input. Conductance streams of 1-20 inputs were then aggregated to implement in whole-cell patch clamp. In a subset of experiments, one spike in the middle of the spike train was synchronized for all individual DCN inputs. In another subset of experiments, pauses of 50 to 800 ms were added to one moment in the spike train to simulate a synchronized pause in DCN activity. To better understand this synapse, a subset of dynamic clamp experiments were done without ISI-dependent scaling (presynaptic plasticity) or NMDA conductances.

To understand how an oscillation in DCN population activity could impact the firing rate of the thalamic neuron, we performed a phase-shift transformation (2) to the *in vivo* DCN input spike train (**Supplemental Figure 5**). The phase-shift transformation was established by two variables: the frequency and amplitude of the baseline oscillation. The frequency varied between 2 Hz or 8 Hz; the amplitude was set to three values: 0.125 (low), 0.25 (medium), and 0.5 (high). Any shifted spike that violated an absolute refractory period of 1ms is dropped. To better characterize this phase shift, we calculated the amplitude of the oscillation in firing rate of the aggregated spike train (10 simulated inputs); for both 2 Hz and 8 Hz trains. The amplitude phase shift resulted in a peak-to-trough ratio of 1.2, medium 1.6, and high 2.7 respectively, and thus represents a physiologically conceivable oscillatory amplitude range of 1.2x to ∼2.5x the baseline firing rate. The conductance streams from this oscillation simulation were then generated as above and used in whole-cell dynamic clamp.

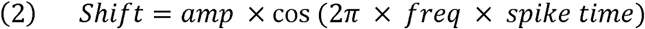

### Viral construction and validation

To generate the viral construct, complementary oligonucleotides encoding the *Cacna1g* gRNA (Forward: 5′-accGGTAGGGCAGCCCCTCCGCCT-3′; Reverse: 5′-aacAGGCGGAGGGGCTGCCCTACC-3′) were annealed and ligated into the SapI site of the pAAV-U6SaCas9gRNA(SapI)-CMV-SaCas9-DIO-pA plasmid. Cultured Neuro2A cells (ATCC CCL-131) were transfected with Lipofectamine 2000 (Thermofisher) using 400 ng of the *Cacna1g* gRNA-containing AAV plasmid or an identical plasmid lacking a gRNA as a negative control. To induce SaCas9 expression, cells were cotransfected with 300 ng of a Cre recombinase expression plasmid (Addgene #201198) and 100 ng of a GFP expression plasmid (Addgene #62916). Seventy-two hours after transfection, cells were resuspended in complete culture medium and subjected to fluorescence-activated cell sorting (FACS) to isolate GFP-positive cells (n = ∼50,000 cells per condition). Genomic DNA was prepared by incubating centrifuged cell pellets in alkaline lysis buffer (25 mM NaOH, 0.2 mM EDTA) at 98°C for 10 min before adding neutralization buffer (40 mM Tris-HCl, pH 5.5).

The genomic region surrounding the target site was PCR amplified using Q5 High-Fidelity DNA Polymerase (New England Biolabs #M0491) and the locus-specific primers Cacna1g_FP (5′-GGAGGGGCTCCGATCGCCCCTCTTCG-3′) and Cacna1g_RP (5′-TGAGGTGTGTGTATTCGGAGGTGGGGCTCG-3′), which generate a 513-bp amplicon encompassing the target site. PCR reactions (50 μL) contained 3 μL of genomic DNA lysate and were amplified using the following cycling conditions: 98°C for 30 s; 40 cycles of 98°C for 10 s, 65°C for 20 s, and 72°C for 20 s; followed by a final extension at 72°C for 2 min. PCR products were purified using silica spin columns (Epoch Life Sciences, #1910-250), and DNA concentrations were determined using a NanoDrop spectrophotometer (Thermofisher).

Genome editing was assessed by restriction fragment length polymorphism (RFLP) analysis. Purified PCR products (200 ng) were digested with 3 U of EciI restriction enzyme in a 15-μL reaction at 37°C for 4 h. In the absence of genome editing, EciI digestion cleaves the 513-bp PCR product into fragments of 302 bp and 211 bp. Genome editing disrupts the EciI recognition sequence, resulting in the persistence of the full-length PCR product. Digested samples were resolved on 2% agarose gels prepared in 0.5X Tris-borate-EDTA (TBE buffer containing 1X SYBR Safe DNA Gel Stain (Invitrogen)). Gels were electrophoresed for approximately 20 min and visualized using a Safe Imager 2.0 Blue-Light Transilluminator (Thermofisher Scientific).

### Behavioral testing

All behavioral testing was performed by an experimenter blind to condition. Prior to training, mice were acclimated to the experimental room for at least 5 hours across 5 days and handled by the experimenter for 5 minutes each day. On training days, mice were acclimated to the testing room for at least 10 minutes. White noise (50-55 dB) was played throughout the animals’ time in the experimental room and light levels were kept stable (650 lux). After histological analysis (below), brains with substantial tissue damage or no evidence of viral expression were excluded from analysis. These determinations were made blind to condition.

### Rotarod

Mice were placed on the experimental apparatus (Pablab Harvard Apparatus LE8205 Rota Rod) and allowed to acclimate to the rod for 15 s before acceleration. The rod accelerated from 4 rpm to 40 rpm over 270 s and rotated for a total of 300 s. The latency to fall was recorded once an animal fell from the rod or stopped running but remained on the rod. Mice were allowed 2 minutes of rest before the subsequent trial. Each animal ran a total of 6 trials per day over 10 days. The apparatus was cleaned with water and unscented soap between groups and at the start and end of day. One mouse was excluded from analysis as it exhibited no learning (<∼15 s on the rotarod every day for the entirety of the 10-day period).

### Gait characterization

Baseline motor behavior was characterized for all mice that underwent behavioral testing using Blackbox Bio automated gait analysis software. On the day prior to recording, the mice were habituated to the Blackbox chamber for the same length of time as the recording itself. The movement of each mouse inside the 30 cm x 30 cm Blackbox chamber was recorded individually for 20 minutes. The Blackbox uses each mouse’s spontaneous movement bouts to calculate gait parameters. Between each mouse, the chamber was cleaned and disinfected with 3% hydrogen peroxide. All Blackbox recordings took place in the same room as behavioral testing above, and the experimenter left the room during the duration of each recording to avoid making noise that might disrupt natural behavior. Lights were kept stable (650 lux) and all mice were kept in their home cages with food and water when not in the chamber and covered with a tarp except for during transfer of mice to and from the chamber.

### Histology

At the conclusion of experiments, animals were anesthetized with isoflurane followed by ketamine/xylazine injection, and then transcardially perfused with 1X PBS (Millipore-Sigma, P3813) then 4% paraformaldehyde (PFA, VWR, 100503-020). The brain was extracted and post-fixed in 4% PFA overnight.

Immunohistochemical protocols were performed to amplify YFP signal for detection in CRISPR-KO animals. Tissue was mounted, immersed in PBS, and sliced into 50 um coronal sections using a vibratome (5100mz, Campden Instruments). For immunohistochemistry, sections were washed with PBS, then incubated for 1 hour with blocking solution containing PBS, 5% normal goat serum (VWR, 101098-382) and 0.2-0.4% TritonX (Millipore-Sigma, T8787). Slices were then immersed in a mixture of blocking solution and primary antibodies (chicken anti-GFP, Abcam ab13970, 1:1000) and incubated overnight at 4°C with agitation. The following day, slices were washed with PBS and incubated in a mixture of blocking solution and secondary antibodies (goat anti-chicken Alexa 488, Abcam ab150173, 1:1000) at room temperature for 1-1.5 hours.

Tissue sections were arranged on microscope slides, then mounted with DAPI Fluoromount solution (Southern Biotech, 0100-20). Fluorescent images of tissue were acquired on a slide scanner microscope (VS200, Evident Scientific).

### Data analysis

All data analysis was performed with custom MATLAB scripts.

### Recording tract reconstruction

Locations of *in vivo* electrophysiology probes were identified post-hoc from 50 um tissue sections sliced and mounted after fixation (see Histological Protocols). Coronal sections containing fluorescent labeling indicative of probe tracts were aligned to representative sections in the Allen Atlas CCFv3^60^. Custom saved trajectories in the Neuropixels Trajectory Explorer (https://github.com/petersaj/neuropixels_trajectory_explorer, by Andy Peters) were used to display the insertion angle, ultimate probe depth, and location of recording channels along the DV recording span. All trajectories are shown on normalized mouse brain with bregma-lambda distance scaled to 4.1 mm.

### Spectral analysis

Recordings were downsampled to 1 kHz. Spectrograms of LFP frequency were generated with a short-time Fourier transform of data, using a Hamming window of 5 s, overlap of 2.5 s, and frequency resolution of 0.12 Hz. Spectrograms displayed power spectral density, and power values were normalized within recordings to “fraction of total power” by dividing by the sum of all values. Dominant frequency was defined as the frequency component with greatest power within each time bin, while preferred dominant frequency was defined as the mode dominant frequency observed within the *delta* band over an entire experiment. *Delta* fraction of power was calculated as the proportion of total power per time bin contained within *delta*-band frequencies. Traditional coherence analysis was not used to determine correlation between LFP signals, since the phase shift between regions created ambiguity about whether *delta* cycles in one region correspond to the preceding or following cycle in another.

Still and move periods were determined from rotary encoder movement speed data, and epochs with a duration of >3 seconds were classified as discrete periods. Power spectra were extracted from individual still/move epochs prior to averaging. Recording periods in which the animal entered NREM or REM sleep were excluded from analysis.

### Phase analysis

Determination of phase of continuous signals was performed by first band-passing the data in the *delta* range (1-5 Hz) and then applying a Hilbert transform to extract instantaneous phase.

For photometrode *delta*-triggered averages, instances when the phase of the DCN axon fluorescence or thalamic LFP signal equaled zero (peak of the *delta* oscillation) were used as triggered timepoints. An identical procedure was conducted for isosbestic control. The shuffled control was computed by taking the *delta* timepoints from each signal, computing the intervals from each timepoint, randomizing the order of those intervals, and then concatenating the new intervals into a shuffled set of timepoints.

For closed-loop stimulation experiments, recording data was filtered using a 2^nd^-order high-pass Butterworth filter for frequencies above 0.7 Hz. For both in-phase and antiphase stimulation recordings, instantaneous *delta*-band phase of thalamic LFP at the time of stimulus was used to classify individual stimuli as rising or falling-phase. To generate baseline STAs, timepoints were extracted at which the filtered trigger signal crossed a voltage threshold empirically determined from stimulation recordings. Timepoints spaced at least 200 ms apart were selected, and in-phase and antiphase conditions were simulated by selecting crossing points with positive or negative slope, respectively. Power spectra represent power spectral density.

### Behavior analysis

Automated machine learning algorithms (Blackbox Bio) were used to automatically generate pose estimation and paw pressure measures. Paw tracking during recording period was used to identify movement bouts, which were defined as at least 3 cycles of continuous steps using all four paws. All gait analysis metrics were performed on bouts then averaged across bouts per animal. Data from sham-injected and empty-vector animals were combined to form the final control group.

## Acknowledgements

This work was supported by grants from the National Institutes of Health: R00NS110978 and R01NS144457 to C.H.C and the generous support of the Grass Foundation. We thank Davide Reato for sharing DCN spike trains and comments. Alex Zhang and Shawn Pavey developed the gait analysis module for the Blackbox. We also thank Court Hull, Jorge Vera Buschmann, Andras Hajnal, Tim Balmer, Wade Regehr, and members of the Chen lab for helpful comments on early drafts of this manuscript.

## REFERENCES

1. Fries, P. Rhythms for Cognition: Communication through Coherence. Neuron 88, 220–235 (2015).

2. Fernandez-Ruiz, A., Sirota, A., Lopes-dos-Santos, V. & Dupret, D. Over and above frequency: Gamma oscillations as units of neural circuit operations. Neuron 111, 936–953 (2023).

3. Vinck, M. et al. Principles of large-scale neural interactions. Neuron 111, 987–1002 (2023).

4. Palmigiano, A., Geisel, T., Wolf, F. & Battaglia, D. Flexible information routing by transient synchrony. Nat. Neurosci. 20, 1014–1022 (2017).

5. Fernández-Ruiz, A. et al. Gamma rhythm communication between entorhinal cortex and dentate gyrus neuronal assemblies. Science 372, eabf3119 (2021).

6. Hutcheon, B., Miura, R. M., Yarom, Y. & Puil, E. Low-threshold calcium current and resonance in thalamic neurons: a model of frequency preference. J. Neurophysiol. 71, 583–594 (1994).

7. Puil, E., Meiri, H. & Yarom, Y. Resonant behavior and frequency preferences of thalamic neurons. J. Neurophysiol. 71, 575–582 (1994).

8. Sherman, S. M. Tonic and burst firing: dual modes of thalamocortical relay. Trends Neurosci. 24, 122–126 (2001).

9. Swadlow, H. A. & Gusev, A. G. The impact of ‘bursting’ thalamic impulses at a neocortical synapse. Nat. Neurosci. 4, 402–408 (2001).

10. Fanselow, E. E., Sameshima, K., Baccala, L. A. & Nicolelis, M. A. L. Thalamic bursting in rats during different awake behavioral states. Proc. Natl. Acad. Sci. 98, 15330–15335 (2001).

11. Borden, P. Y. et al. Thalamic bursting and the role of timing and synchrony in thalamocortical signaling in the awake mouse. Neuron 110, 2836–2853.e8 (2022).

12. Jahnsen, H. Electrophysiological characteristics of neurones in the guinea-pig deep cerebellar nuclei in vitro. J. Physiol. 372, 129–147 (1986).

13. Reato, D., Tara, E. & Khodakhah, K. Deep Cerebellar Nuclei Rebound Firing In Vivo. in The Neuronal Codes of the Cerebellum 27–51 (Elsevier, 2016). doi:10.1016/B978-0-12-801386-1.00002-2.

14. Aizenman, C. D. & Linden, D. J. Regulation of the Rebound Depolarization and Spontaneous Firing Patterns of Deep Nuclear Neurons in Slices of Rat Cerebellum. J. Neurophysiol. 82, 1697– 1709 (1999).

15. Gornati, S. V. et al. Differentiating Cerebellar Impact on Thalamic Nuclei. Cell Rep. 23, 2690–2704 (2018).

16. Liu, C.-W. et al. The cerebellum shapes motions by encoding motor frequencies with precision and cross-individual uniformity. *Nat*. Biomed. Eng. 9, 1952–1971 (2025).

17. Wang, Y.-M. et al. Neuronal population activity in the olivocerebellum encodes the frequency of essential tremor in mice and patients. Sci. Transl. Med. 16, eadl1408 (2024).

18. Hamel-Thibault, A., Thénault, F., Whittingstall, K. & Bernier, P.-M. Delta-Band Oscillations in Motor Regions Predict Hand Selection for Reaching. Cereb. Cortex cercor;bhw392v1 (2016) doi:10.1093/cercor/bhw392.

19. Spampinato, D. A., Antonioni, A., D’Angelo, E. & Koch, G. Cerebellar rhythms: mechanisms, functions and translational opportunities. Nat. Rev. Neurosci. 10.1038/s41583-026-01072-y (2026) doi:10.1038/s41583-026-01072-y.

20. Okun, M., Naim, A. & Lampl, I. The Subthreshold Relation between Cortical Local Field Potential and Neuronal Firing Unveiled by Intracellular Recordings in Awake Rats. J. Neurosci. 30, 4440– 4448 (2010).

21. Steriade, M., Nunez, A. & Amzica, F. Intracellular analysis of relations between the slow (& 1 Hz) neocortical oscillation and other sleep rhythms of the electroencephalogram. J. Neurosci. 13, 3266–3283 (1993).

22. Contreras, D. & Steriade, M. Cellular basis of EEG slow rhythms: a study of dynamic corticothalamic relationships. J. Neurosci. 15, 604–622 (1995).

23. Kebschull, J. M. et al. Cerebellar nuclei evolved by repeatedly duplicating a conserved cell-type set. Science 370, eabd5059 (2020).

24. Fujita, H., Kodama, T. & Du Lac, S. Modular output circuits of the fastigial nucleus for diverse motor and nonmotor functions of the cerebellar vermis. eLife 9, e58613 (2020).

25. Kang, S. et al. Recent Advances in the Understanding of Specific Efferent Pathways Emerging From the Cerebellum. Front. Neuroanat. 15, 759948 (2021).

26. Chen, C., Blitz, D. M. & Regehr, W. G. Contributions of Receptor Desensitization and Saturation to Plasticity at the Retinogeniculate Synapse. Neuron 33, 779–788 (2002).

27. Schäfer, C. B., Gao, Z., De Zeeuw, C. I. & Hoebeek, F. E. Temporal dynamics of the cerebello-cortical convergence in ventro-lateral motor thalamus. J. Physiol. 599, 2055–2073 (2021).

28. Chen, C. & Regehr, W. G. Developmental Remodeling of the Retinogeniculate Synapse. Neuron 28, 955–966 (2000).

29. Koster, K. P. & Sherman, S. M. Convergence of inputs from the basal ganglia with layer 5 of motor cortex and cerebellum in mouse motor thalamus. eLife 13, e97489 (2024).

30. Canto, C. B., Witter, L. & De Zeeuw, C. I. Whole-Cell Properties of Cerebellar Nuclei Neurons In Vivo. PLOS ONE 11, e0165887 (2016).

31. Fogerson, P. M. & Huguenard, J. R. Tapping the Brakes: Cellular and Synaptic Mechanisms that Regulate Thalamic Oscillations. Neuron 92, 687–704 (2016).

32. Contreras, D., Destexhe, A., Sejnowski, T. J. & Steriade, M. Control of Spatiotemporal Coherence of a Thalamic Oscillation by Corticothalamic Feedback. Science 274, 771–774 (1996).

33. Zingg, B. et al. AAV-Mediated Anterograde Transsynaptic Tagging: Mapping Corticocollicular Input-Defined Neural Pathways for Defense Behaviors. Neuron 93, 33–47 (2017).

34. Hunnicutt, B. J. et al. A comprehensive thalamocortical projection map at the mesoscopic level. Nat. Neurosci. 17, 1276–1285 (2014).

35. Kakei, S., Na, J. & Shinoda, Y. Thalamic terminal morphology and distribution of single corticothalamic axons originating from layers 5 and 6 of the cat motor cortex. J. Comp. Neurol. 437, 170–185 (2001).

36. Na, J., Kakei, S. & Shinoda, Y. Cerebellar input to corticothalamic neurons in layers V and VI in the motor cortex. Neurosci. Res. 28, 77–91 (1997).

37. Roth, R. H. et al. Thalamic integration of basal ganglia and cerebellar circuits during motor learning. Preprint at 10.1101/2024.10.31.621388 (2024).

38. Varani, A. P. et al. Multiple functions of cerebello-thalamic neurons in learning and offline consolidation of a motor skill in mice. eLife 13, RP102813 (2026).

39. Sala, R. W., Ayyaz, A., Léna, C. & Popa, D. Offline cerebello-cortico-striatal dynamics predict motor strategy exploration and retention in skill learning. Preprint at 10.64898/2025.12.09.693142 (2025).

40. Lee, J.-H. et al. Cerebellar granule cell signaling is indispensable for normal motor performance. Cell Rep. 42, 112429 (2023).

41. Antal, M., Beneduce, B. M. & Regehr, W. G. The Substantia Nigra Conveys Target-Dependent Excitatory and Inhibitory Outputs from the Basal Ganglia to the Thalamus. J. Neurosci. 34, 8032– 8042 (2014).

42. Crandall, S. R., Cruikshank, S. J. & Connors, B. W. A Corticothalamic Switch: Controlling the Thalamus with Dynamic Synapses. Neuron 86, 768–782 (2015).

43. Martinez-Garcia, R. I. et al. Two dynamically distinct circuits drive inhibition in the sensory thalamus. Nature 583, 813–818 (2020).

44. Lam, Y.-W. & Sherman, S. M. Functional topographic organization of the motor reticulothalamic pathway. J. Neurophysiol. 113, 3090–3097 (2015).

45. Cox, C. L., Huguenard, J. R. & Prince, D. A. Nucleus reticularis neurons mediate diverse inhibitory effects in thalamus. Proc. Natl. Acad. Sci. 94, 8854–8859 (1997).

46. Molineux, M. L. et al. Specific T-type calcium channel isoforms are associated with distinct burst phenotypes in deep cerebellar nuclear neurons. Proc. Natl. Acad. Sci. 103, 5555–5560 (2006).

47. Yunker, A. M. R. et al. Immunological characterization of T-type voltage-dependent calcium channel CaV3.1 (alpha1G) and CaV3.3 (alpha1I) isoforms reveal differences in their localization, expression, and neural development. Neuroscience 117, 321–335 (2003).

48. Matsumoto-Makidono, Y. et al. Ionic Basis for Membrane Potential Resonance in Neurons of the Inferior Olive. Cell Rep. 16, 994–1004 (2016).

49. Khosrovani, S., Van Der Giessen, R. S., De Zeeuw, C. I. & De Jeu, M. T. G. *In vivo* mouse inferior olive neurons exhibit heterogeneous subthreshold oscillations and spiking patterns. Proc. Natl. Acad. Sci. 104, 15911–15916 (2007).

50. Coffman, K. A., Dum, R. P. & Strick, P. L. Cerebellar vermis is a target of projections from the motor areas in the cerebral cortex. Proc. Natl. Acad. Sci. 108, 16068–16073 (2011).

51. Suzuki, L., Coulon, P., Sabel-Goedknegt, E. H. & Ruigrok, T. J. H. Organization of Cerebral Projections to Identified Cerebellar Zones in the Posterior Cerebellum of the Rat. J. Neurosci. 32, 10854–10869 (2012).

52. Desai, N. V. & Varela, C. Distinct burst properties contribute to the functional diversity of thalamic nuclei. J. Comp. Neurol. 529, 3726–3750 (2021).

53. Pisano, T. J. et al. Homologous organization of cerebellar pathways to sensory, motor, and associative forebrain. Cell Rep. 36, 109721 (2021).

54. Margrie, T., Brecht, M. & Sakmann, B. In vivo, low-resistance, whole-cell recordings from neurons in the anaesthetized and awake mammalian brain. Pfl gers Arch. Eur. J. Physiol. 444, 491–498 (2002).

55. Lopes, G. et al. Bonsai: an event-based framework for processing and controlling data streams. Front. Neuroinformatics 9, (2015).

56. Turecek, J., Jackman, S. L. & Regehr, W. G. Synaptotagmin 7 confers frequency invariance onto specialized depressing synapses. Nature 551, 503–506 (2017).

57. Marino, M., Misuri, L. & Brogioli, D. A new open source software for the calculation of the liquid junction potential between two solutions according to the stationary Nernst-Planck equation. Preprint at 10.48550/ARXIV.1403.3640 (2014).

58. Dolzer, J. Patch Clamp Technology in the Twenty-First Century. in Patch Clamp Electrophysiology (eds Dallas, M. & Bell, D.) vol. 2188 21–49 (Springer US, New York, NY, 2021).

59. Jahr, C. & Stevens, C. Voltage dependence of NMDA-activated macroscopic conductances predicted by single-channel kinetics. J. Neurosci. 10, 3178–3182 (1990).

60. Wang, Q. et al. The Allen Mouse Brain Common Coordinate Framework: A 3D Reference Atlas. Cell 181, 936–953.e20 (2020).

