## Supplemental Data for "The cerebellum exploits a thalamic carrier frequency to refine motor behaviors"

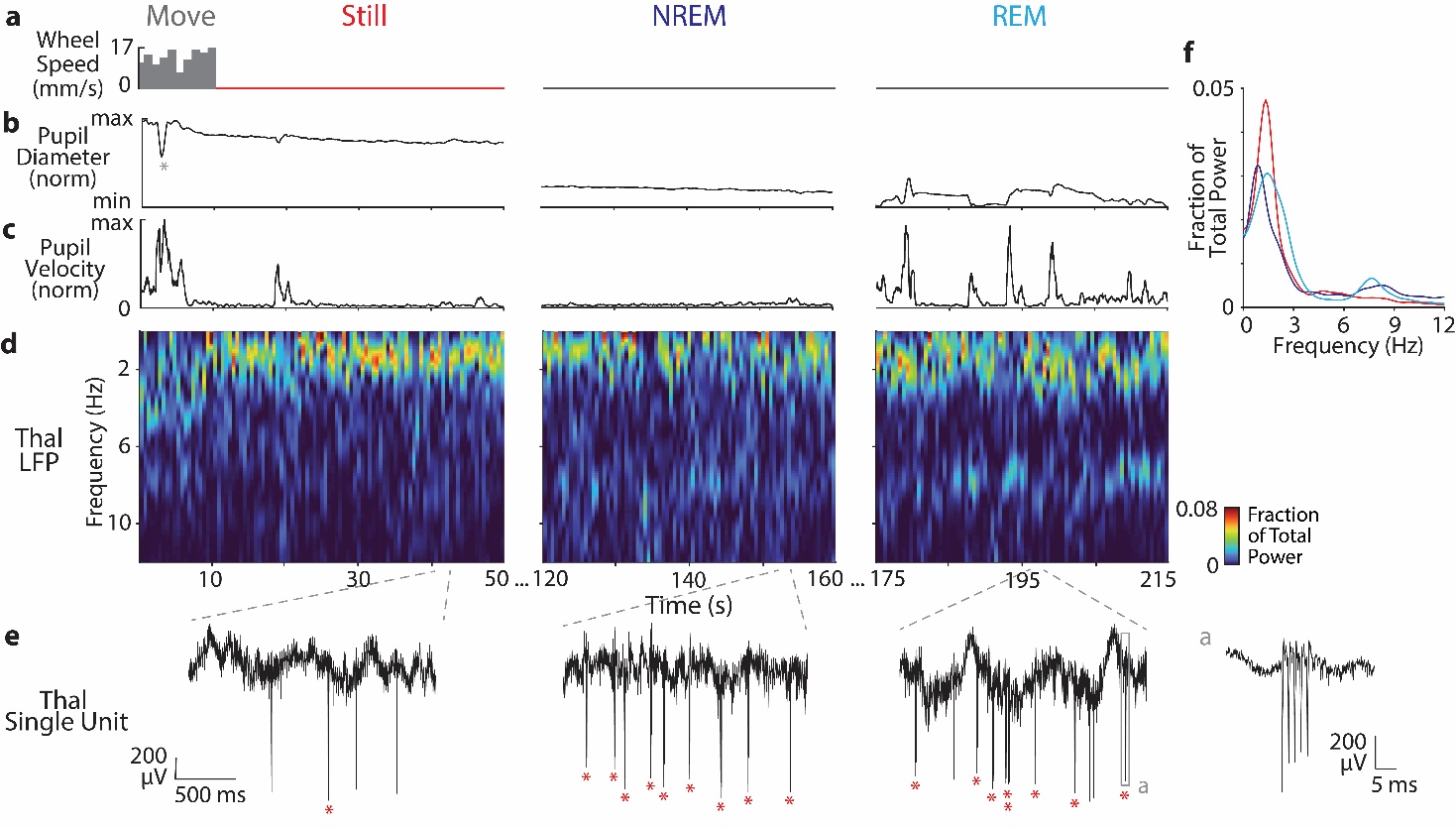

**Supplemental Figure 1. Awake, still-period cerebellothalamocortical *delta* oscillations are dissociable from sleep. (A)** Wheel speed indicating movement during epochs of movement, stillness, NREM and REM sleep. **(B)** Normalized pupil diameter through all behavioral states. Pupil constricts to a high degree during sleep. Gray asterisk denotes an eyeblink. **(C)** Pupil velocity through all arousal states. Periods of high velocity in REM sleep represent saccadic eye movements. **(D)** Representative spectrograms of thalamic LFP in all behavioral states. **(E)** Activity of a sample thalamic single unit tracked through awake stillness, NREM and REM sleep. Red asterisks denote bursts. **(Ea)** An example burst. **(F)** Power spectral density (PSD) plots of thalamic LFP by arousal state. Thalamic *delta* fraction of power in awake, still epochs can exceed NREM and REM.

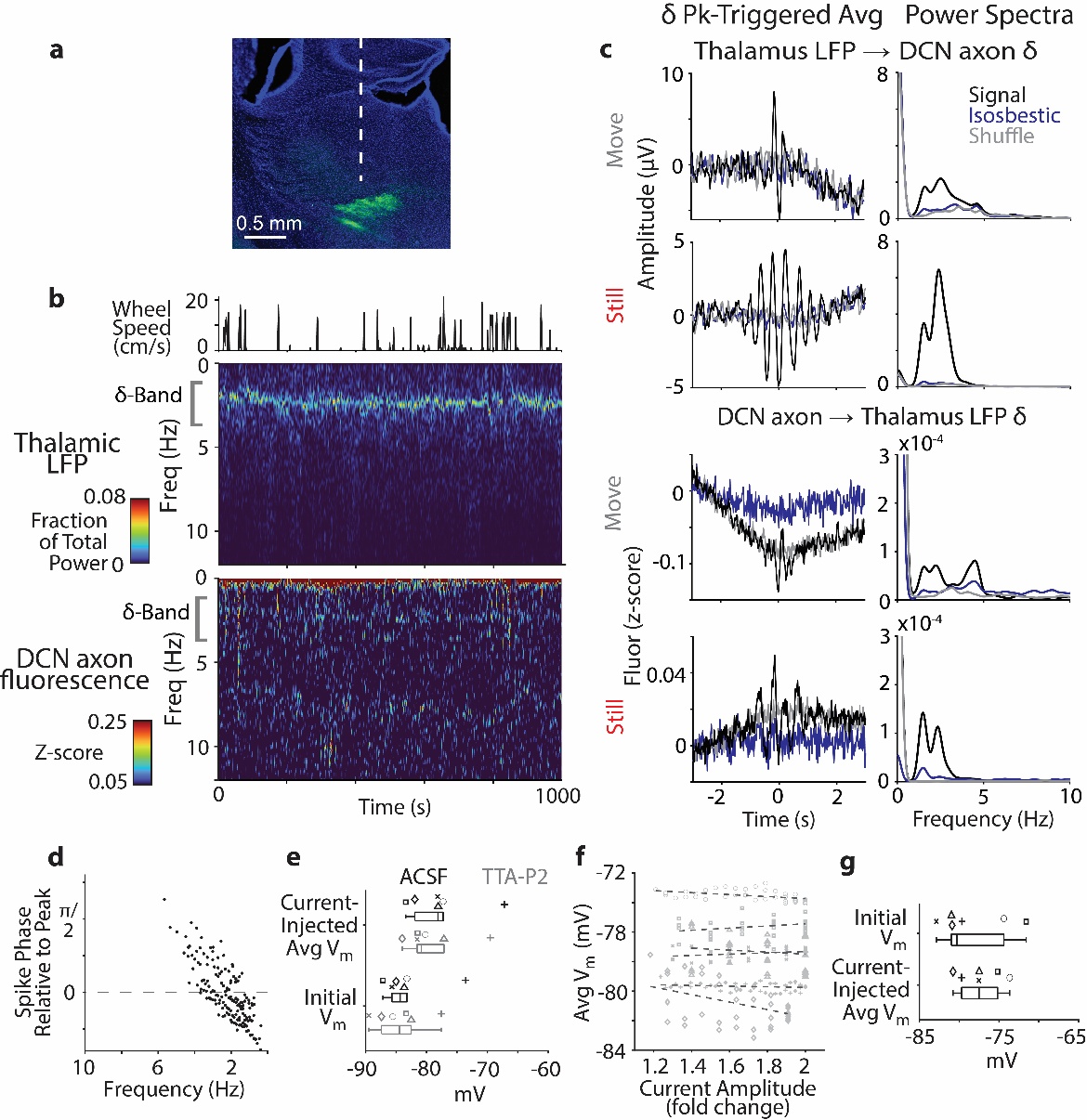
**Supplemental Figure 2. Frequency-based cerebellothalamic transmission depends on CaV3.1 dynamics. (A)** Example photometrode trajectory (dashed line) in thalamus. DCN axon terminals seen in green. **(B)** Example photometrode recording, showing movement bouts (top) with spectrograms of thalamic LFP (middle) and DCN axon fluorescence signal (bottom). Note concurrent increase in power in the *delta* bands. **(C)** Comparison of *delta* peak-triggered averages (left column) and PSDs (right column) in move and still periods between experimental thalamic LFP and DCN axon signals, isosbestic channels, and peak-timepoint-shuffled control. **(D)** Per-cycle latency between phase of spike response and current-injection peak in current-clamp chirp experiments. Spikes occur on rising phase of current oscillation within *delta* band and within the falling phase at higher frequencies. **(E)** Cell membrane potentials prior to (“initial”) and during current injection, before and after TTA-P2 wash-on. Symbols represent individual cells. Similar values between cells in ACSF and TTA-P2 experiments attributes changes in burst response to CaV3.1. **(F)** Average membrane potential per cycle for current-clamp amplitude modulation experiments (markers denote single cycles, lines indicate linear fit to single cell). **(G)** Cell membrane potentials prior to and during current injection. Symbols denote individual cells. Similar values between experimental cells attributes burst response to changes in current amplitude.

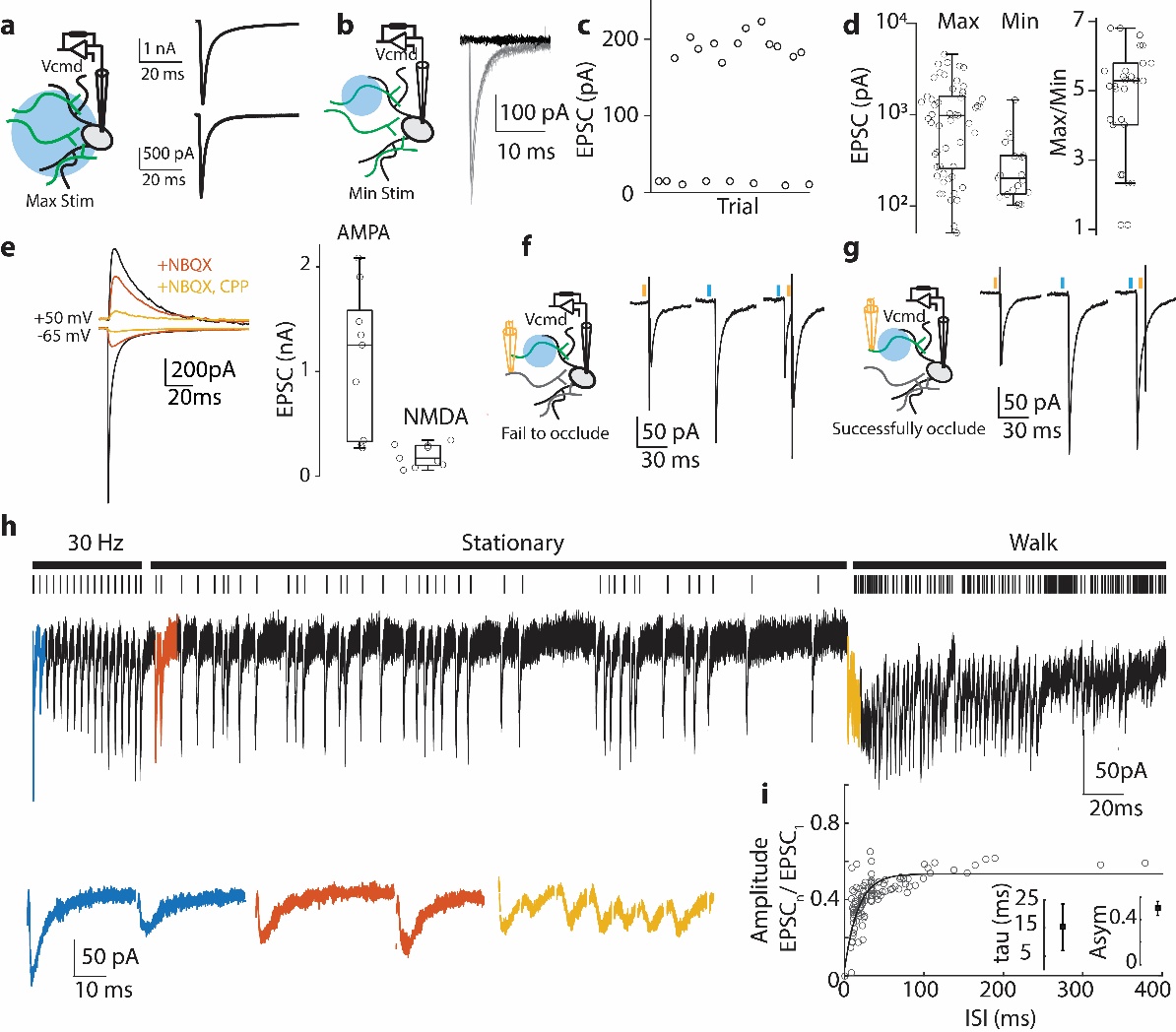

**Supplemental Figure 3. Characterization of cerebellar inputs to the VAL for dynamic clamp**. ChR2 was expressed in DCN axons, and coronal slices of the thalamus were made. **(A)** Left: whole-cell recordings were made in thalamic neurons held at -65 mV, and full-field optogenetic stimuli (473 nm) were delivered. Right: two example responses. **(B)** Light was constrained to a ~70 μm spot size to conduct minimal stimulation experiments. Right: example experiment with responses overlaid. **(C)** Quantification of evoked response amplitude of the example shown in (**B**). **(D)** Full-field (max) and minimal (min) evoked current (left) and the average max/min current ratio (right), which approximates the number of inputs per cell. **(E)** Isolated AMPA and NMDA components. Left: example currents. Right: amplitude of evoked currents at -65 mV. **(F)** Optogenetic occlusion^56^ was used to isolate cerebellar inputs for electrical stimulation. Example of a failed occlusion. Orange ticks indicate electrical stimulus timing; blue optogenetic. **(G)** Example of a successful occlusion. **(H)** Isolated inputs were given a 30-Hz stimulus (the average baseline firing rate of DCN neurons) followed by “still” and “move” spike patterns as measured *in vivo*^13^. Top raster indicates stimulus timing, evoked currents on the bottom (stim artifacts removed, enlarged currents correspond as per indicated colors). **(I)** Amount of depression observed for each interstimulus interval (ISI). Inset: averaged tau and steady-state depression values (asym) across all experiments (n = 4).

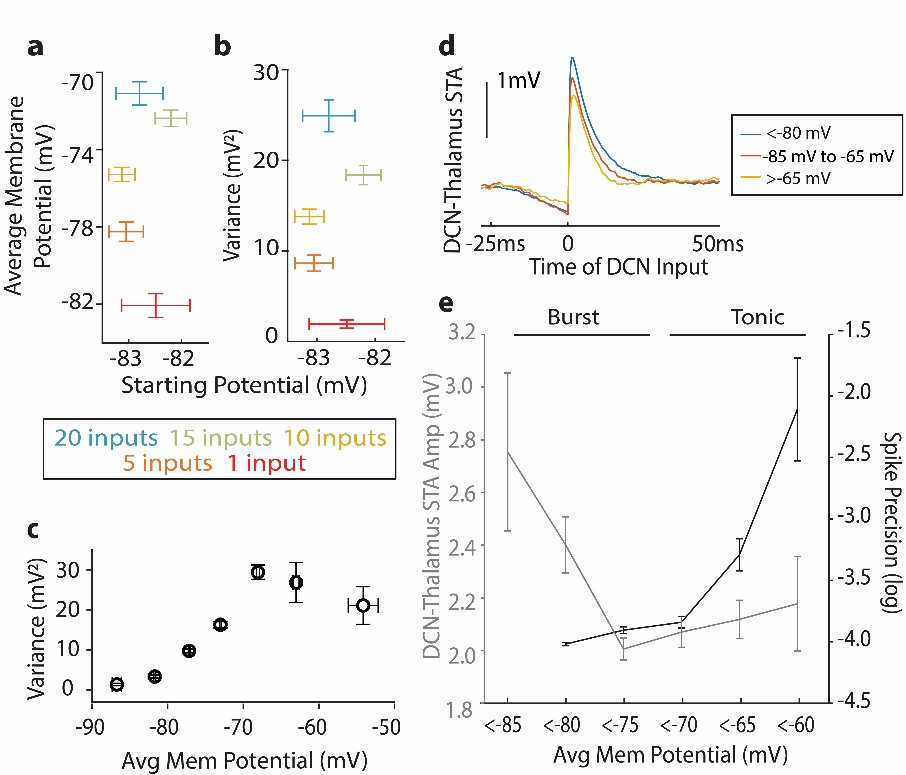

**Supplemental Figure 4.** **Dynamic clamp matches parameters measured *in vivo*. (A)** Varying numbers of inputs were used to investigate the range of thalamic response to DCN inputs. Cells were held at similar starting potentials before conductances were injected (n = 67). The average membrane potential in response to conductance streams was calculated for all trials across numbers of input. Higher input numbers results in more depolarized potentials. **(B)** Variance of membrane potential is similarly calculated. Increased variance is also found with increased inputs. Notably, variance for 10-15 inputs most resembles variance seen in *in vivo* experiments (**Figure 1G**). **(C)** Trials were further sorted by their membrane potential during conductance injection and variance was found to vary in a nonlinear fashion. The variance peaks at trials with membrane potentials of -70 mV, which most resembles physiological potential maintained in burst mode *in vivo* (**Figure 1F**). Trials more depolarized than -70 mV had lower variance. **(D)** DCN input-triggered average shows membrane potential-dependent differences in the EPSP. **(E)** Quantification of DCN input-triggered average and spike precision (multiplicative inverse of iFR standard deviation) value of trials binned by membrane potential showed increased EPSP amplitudes in burst mode and greater spike precision in tonic.

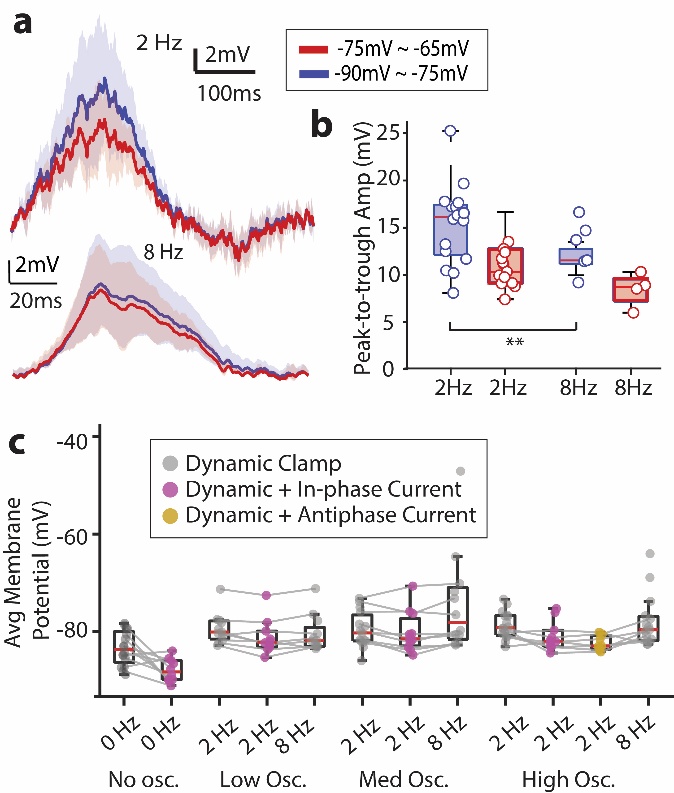

**Supplemental Figure 5: *Delta*-oscillating DCN input enhances low-threshold bursts. (A)** When sorted by the trial-averaged membrane potential, oscillation cycle-averaged membrane potential showed a boost in the burst mode trials when compared to tonic mode trials only in the 2-Hz condition. Trials with additional oscillatory current were excluded from this analysis. **(B)** Cell-matched averaged quantification of the peak-to-trough difference (from **A**) in membrane potential shows significant increase in amplitude in burst mode with 2 Hz oscillation. 8 Hz oscillation of peak to trough amplitude shows no difference in membrane potential amplitude. **(C)** Quantification of membrane potential in response to varying frequency, oscillation amplitude, and current injection (n = 18).

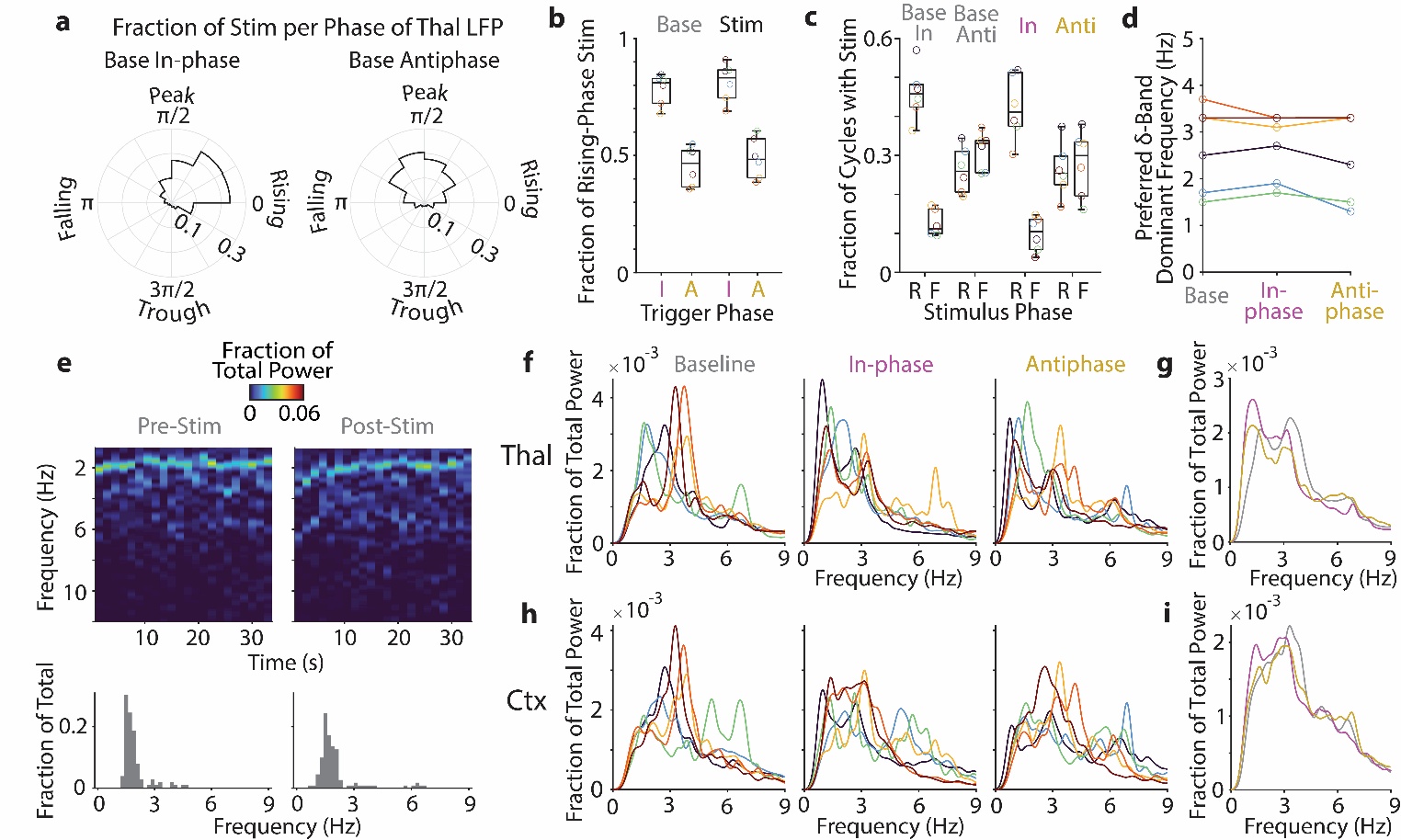

**Supplemental Figure 6. Closed-loop in-phase and antiphase DCN stimulation.** For accurate characterization of LFP response to in-phase and antiphase stimulation (**Figure 4J**), timepoints of electrical stimulation were simulated for no-stimulus baseline recordings. **(A)** Distribution of simulated stimulation timepoints among thalamic *delta*-band phase in baseline recordings. Simulated timepoints for in-phase and antiphase conditions were extracted by applying voltage threshold and refractory period criteria to baseline recordings (see **Methods**). Stimuli with instantaneous phase from trough to peak were classified as “rising-phase” with those during peak to trough as “falling-phase”. Stimulation affecting intrinsic thalamic *delta* phase precluded the ability to interpret this analysis for real stimuli. **(B-C)** We validated the baseline simulation with respect to stimulus conditions by comparing the number of evoked stimuli with respect to phase. **(B)** Stimulation accuracy for simulated vs. real stimuli, represented as proportion of rising-phase stimuli during in- vs. antiphase stimulation (baseline in-phase median: 0.811, baseline antiphase median: 0.467, stim in-phase median: 0.832, stim antiphase median: 0.484). Note similar values between baseline and stimulation categories. **(C)** Stimulation consistency for simulated vs. real stimuli, represented as number of *delta*-band cycles containing rising- and falling-phase stimuli as a fraction of total cycle number (R = rising-phase, F = falling-phase). In-phase recordings displayed a greater fraction of rising-phase stimuli, whereas antiphase recordings had a relatively greater preference for falling-phase stimuli. **(D)** The preferred dominant frequency within the *delta* band for baseline recordings was preserved during stimulation. **(E)** Example spectrograms (top) and dominant frequency histograms (bottom) for baseline recordings made before and after stimulation recordings, depicting the transient effects of stimulation on *delta*-band activity. **(F)** PSDs of thalamic LFP by individual recording (n = 6). Colors for individual experiments match those in (**B-D**) and **Figure 4F-H**. **(G)** Thalamic PSDs in (**F**) averaged between recordings for baseline (gray), in-phase (magenta) and antiphase (gold) conditions. **(H-I)** Same as (**F-G**), for cortical LFPs.

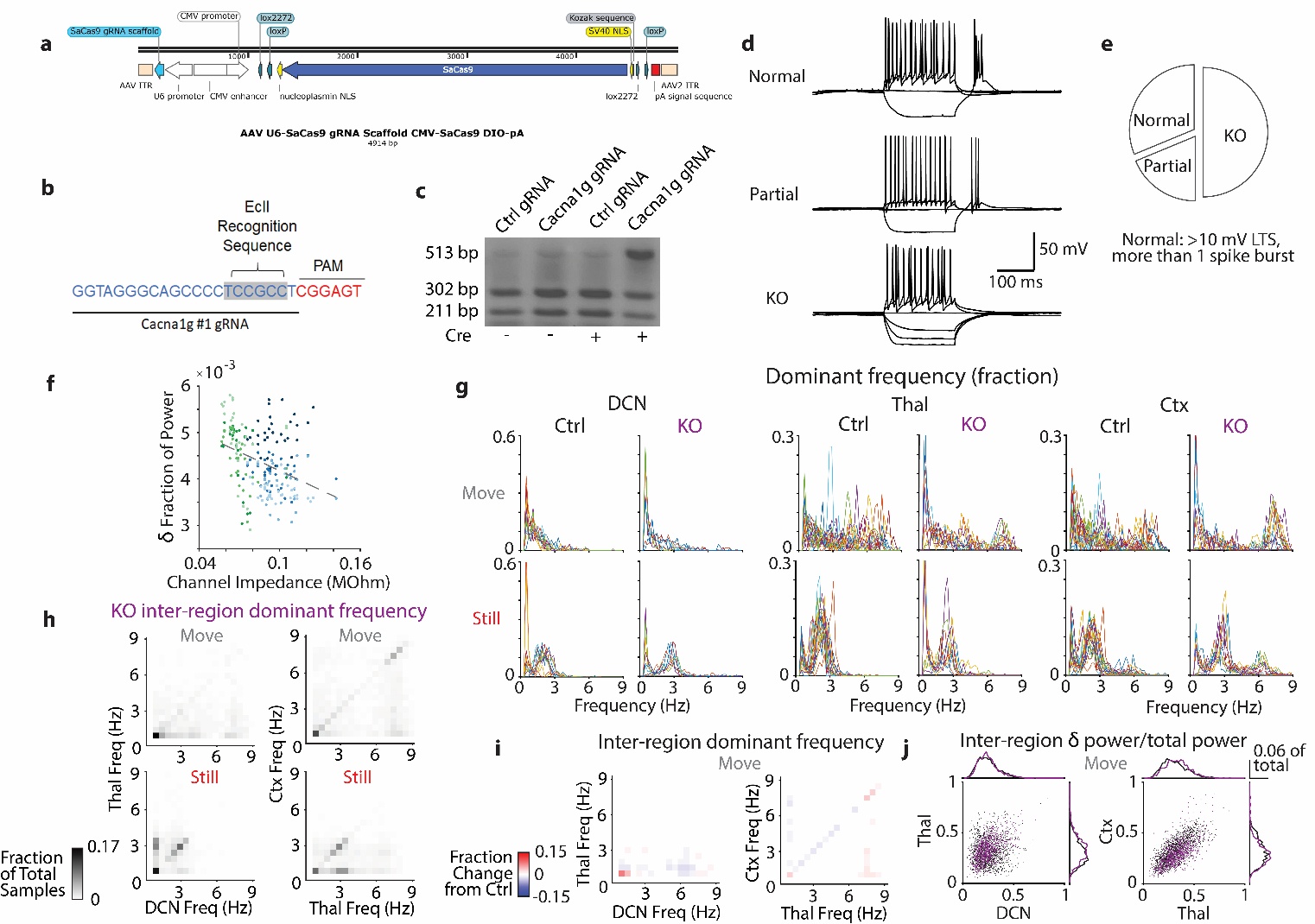
**Supplemental Figure 7: Validation of CRISPR-KO phenotype. (A)** AAV vector construct for expression of U6-promoter gRNA and CMV-promoter DIO SaCas9. **(B)** Blue: Cacna1g gRNA sequence; red: associated SaCas9 PAM sequence. Excision of EciI recognition sequence blocks restriction enzyme cleavage, enabling indel detection. **(C)** Stained gel electrophoresis results for Cre- and viral plasmid-transfected Neuro2a cells following PCR amplification of locus in (**B**). 513-bp band indicates increased indel occurrence and successful knockout only in presence of Cre and *Cacna1g* gRNA. **(D)** *In vitro* electrophysiological validation of CaV3.1 knockout. Motor thalamic cells exhibited normal burst response (top), partial knockout (middle, diminished LTS and/or limited rebound firing), or total knockout (bottom, no rebound response). **(E)** Proportion of total cells (n = 16) exhibiting each knockout phenotype. **(F)** To confirm that the location dependence of *delta* power (**Figure 5D**) does not instead result from differences in probe channel impedance, we calculated the correlation between impedance and *delta* fraction of power. Dotted line represents linear model fit (R-sq = 0.142, note low value). **(G)** PSDs of move vs. still periods by region for control vs. KO animals. Line colors represent individual periods. **(H)** Inter-region dominant-frequency correlations as in **Figure 1M**, for KO animals. Note diminished fraction of total samples in still-period *delta* band. **(I)** Subtracted dominant frequency heatmaps as in **Figure 5G**, for move periods. **(J)** Inter-region *delta* fraction of power as in **Figure 5H**, in move periods.

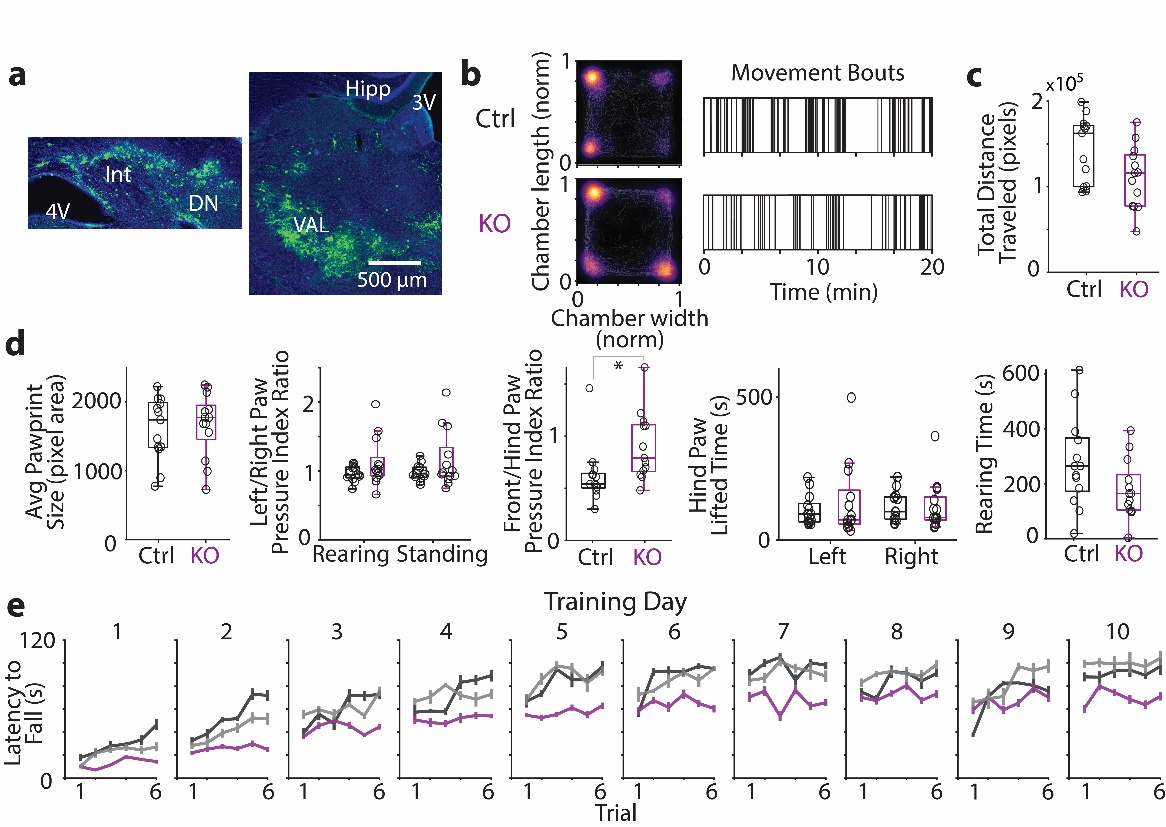
**Supplemental Figure 8: Baseline movement kinematics are preserved in CaV3.1 knockout animals. (A)** Left: DCN tissue from KO animal infected with AAV-Cre and coinjected with a Cre-dependent YFP virus (green). Right: VAL tissue from KO animal coinjected with AAV-CRISPR-saCas9 and a Cre-dependent YFP virus (green). **(B)** Left: Normalized position heatmaps of example control and KO mouse within the Blackbox chamber for a 20-minute recording. Right: Raster plots of movement bout distribution during recording. **(C)** Total pixel distance traveled in chamber for combined control (N = 13) vs. KO (N = 13) groups (p = 0.1731). All statistical analysis for gait parameters used the FDR-corrected Kruskal-Wallis test. **(D)** Blackbox gait parameters remain comparable between control and KO groups. Left to right: Average pawprint area in pixels (p = 0.8530). Paw pressure ratio between left and right paws (rearing: p = 0.6623, standing: p = 0.5336). Paw pressure ratio between front and hind paws (p = 0.0454). Total recording time where hind paws are lifted (left: p = 0.8530, right: p = 0.6624). Total rearing time (both front paws lifted; p = 0.2574). **(E)** Intra-day latency-to-fall trajectories for sham-injected (black, N = 6), empty-vector (gray, N = 7) and KO (violet, N = 13) mice on days 1-10 of accelerating rotarod task. Sham and empty-vector groups were combined into a final control group for **B-D** and **Figure 6**. Note that on Day 9, construction on the floor below the lab elicited loud drilling noises that reverberated throughout our behavior room, resulting in an uncharacteristic dip in performance in a cohort of animals.

| **Figure** | **Test** | **Comparison** | **Test Stats** | **P-value** | **Signif?** |
| --- | --- | --- | --- | --- | --- |
| Fig 1L (averages) | Unpaired Wilcoxon rank sum, two-tailed | DN Dominant Frequency Move vs. Still | ranksum: 2.5202e06, zstat: -18.8144 | 5.75e-79 | X |
|  | Unpaired Wilcoxon rank sum, two-tailed | Thal Dominant Frequency Move vs. Still | ranksum: 1.0065e07, zstat: 19.3779 | 1.19e-83 | X |
|  | Unpaired Wilcoxon rank sum, two-tailed | Ctx Dominant Frequency Move vs. Still | ranksum: 9.1243e06, zstat: 8.7739 | 1.73e-18 | X |
| Fig 1N (histograms) | Unpaired Wilcoxon rank sum, one-tailed | DN Ctrl Move Power vs. DN Ctrl Still Power | ranksum: 919986, zstat: -22.7895, Cohen’s *d*: -0.6378 | 2.91e-115 | X |
|  | Unpaired Wilcoxon rank sum, one-tailed | Thal Ctrl Move Power vs. Thal Ctrl Still Power | ranksum: 804341, zstat: -27.7232, Cohen’s *d*: -2.4114 | 1.83e-169 | X |
|  | Unpaired Wilcoxon rank sum, one-tailed | Ctx Ctrl Move Power vs. Ctx Ctrl Still Power | ranksum: 867366, zstat: -25.0344, Cohen’s *d*: -1.4273 | 1.29e-138 | X |
| Fig 2E | Linear model goodness-of-fit | DCN dominant frequency vs. Thal dominant frequency | R-sq: 0.996 | 2.56e-05 | X |
| Fig 2F | Unpaired Wilcoxon rank sum, one-tailed | DCN🡪Thal, Ca vs. 1 | ranksum: 51 | 0.0238 | X |
|  |  | DCN🡪Thal, Isos vs. 1 | ranksum: 39 | 0.5498 |  |
|  |  | DCN🡪Thal, Shuff vs. 1 | ranksum: 27 | 0.9935 |  |
|  | Unpaired Wilcoxon rank sum, one-tailed | Thal🡪DCN, Ca vs. 1 | ranksum: 51 | 0.0011 | X |
|  |  | Thal🡪DCN, Isos vs. 1 | ranksum: 21 | 1 |  |
|  |  | Thal🡪DCN, Shuff vs. 1 | ranksum: 27 | 0.9935 |  |
| Supp Fig 2E | Paired Wilcoxon signed rank, two-tailed | ACSF vs. TTA current-injection Vm | signedrank: 6 | 0.4375 |  |
|  | Paired Wilcoxon signed rank, two-tailed | ACSF vs. TTA initial Vm | signedrank: 3 | 0.1562 |  |
| Fig 3J Spike Rates | Paired, Wilcoxon signed ranks, one-tailed | 2Hz vs 8Hz at low amp | N = 10; Rank sum = 53 | 0.0029 | X |
|  | Paired, Wilcoxon signed ranks, one-tailed | 2Hz vs 8Hz at med amp | N = 12; Rank sum = 66 | 4.88E-04 | X |
|  | Paired, Wilcoxon signed ranks, one-tailed | 2Hz vs 8Hz at high amp | N = 15; Rank sum = 120 | 3.05E-05 | X |
| Fig 3K Spikes with iFR > 100 | Paired, Wilcoxon signed ranks, one-tailed | 2Hz vs 8Hz at low amp | N = 6; Rank sum = 21 | 0.016 | X |
|  | Paired, Wilcoxon signed ranks, one-tailed | 2Hz vs 8Hz at med amp | N = 7; Rank sum = 28 | 0.0078 | X |
|  | Paired, Wilcoxon signed ranks, one-tailed | 2Hz vs 8Hz at high amp | N = 12; Rank sum = 66 | 4.88E-04 | X |
| Fig 4C Spike rates | Paired, Wilcoxon signed ranks, one-tailed | 2Hz high amp: standard vs with oscillating current | N = 10; Rank sum = 16.5 | 0.14 |  |
|  | Paired, Wilcoxon signed ranks, one-tailed | 2Hz high amp: standard vs with oscillating antiphase current | N = 7; Rank sum = 28 | 0.0078 | X |
|  | Paired, Wilcoxon signed ranks, one-tailed | 2hz high amp: inphase current vs antiphase current | N = 7; Rank sum = 28 | 0.0078 | X |
| Fig 4C Spikes with iFR > 100 | Paired, Wilcoxon signed ranks, one-tailed | 2Hz high amp: standard vs with oscillating current | N = 9; Rank sum = 19 | 0.36 |  |
|  | Paired, Wilcoxon signed ranks, one-tailed | 2Hz high amp: standard vs with oscillating antiphase current | N = 5; Rank sum = 15 | 0.031 | X |
|  | Paired, Wilcoxon signed ranks, one-tailed | 2Hz high amp: inphase vs antiphase current | N = 6; Rank sum = 21 | 0.015 | X |
| Fig 4F | Paired, Wilcoxon signed ranks, one-tailed | Mean Prob of Delta Dom Freq, In-phase vs. 1 | N = 7; Rank sum = 28 | 0.0078 | X |
|  | Paired, Wilcoxon signed ranks, one-tailed | Mean Prob of Delta Dom Freq, Antiphase vs. 1 | N = 7; Rank sum = 0 | 0.0078 | X |
| Fig 4G | Paired, Wilcoxon signed ranks, one-tailed | PSD Delta AUC, In-phase vs. 1 | N = 7; Rank sum = 27 | 0.016 | X |
|  | Paired, Wilcoxon signed ranks, one-tailed | PSD Delta AUC, Antiphase vs. 1 | N = 7; Rank sum = 0 | 0.0078 | X |
| Fig 4M | Unpaired Wilcoxon rank sum, one-tailed | Mean Prob of Delta Dom Freq, In-phase vs. 1 | ranksum: 51 | 0.9935 |  |
|  | Unpaired Wilcoxon rank sum, one-tailed | Mean Prob of Delta Dom Freq, Antiphase vs. 1 | ranksum: 57 | 0.0011 | X |
| Fig 4N | Unpaired Wilcoxon rank sum, one-tailed | PSD Delta AUC, In-phase vs. 1 | ranksum: 45 | 0.8680 |  |
|  | Unpaired Wilcoxon rank sum, one-tailed | PSD Delta AUC, Antiphase vs. 1 | ranksum: 51 | 0.0238 | X |
| Supplemental Fig 5F Hyperpolarized (-90~-75 mV) group | Paired, Wilcoxon signed ranks, two-tailed | 2hz vs 8Hz | N = 13; Rank sum = 88 | 0.0012 | X |
| Depolarized (-75~ -65 mV) group | Paired, Wilcoxon signed ranks, two-tailed | 2hz vs 8Hz | N = 3; Rank sum = 6 | 0.25 |  |
| Supplemental Fig 5C Membrane Potential | Paired, Wilcoxon signed ranks, two-tailed | 2Hz vs 8Hz at low amp | N = 12; ranksum: 30 | 0.52 |  |
|  | Paired, Wilcoxon signed ranks, two-tailed | 2Hz vs 8Hz at med amp | N = 15; ranksum: 57 | 0.89 |  |
|  | Paired, Wilcoxon signed ranks, two-tailed | 2Hz vs 8Hz at high amp | N = 10; ranksum 43 | 0.13 |  |
|  | Paired, Wilcoxon signed ranks, two-tailed | 2Hz high amp: standard vs with oscillating current | N = 10; Rank sum = 16.5 | 0.29 |  |
|  | Paired, Wilcoxon signed ranks, two-tailed | 2Hz high amp: standard vs with oscillating antiphase current | N = 7; Rank sum = 28 | 0.017 | X |
|  | Paired, Wilcoxon signed ranks, two-tailed | 2hz high amp: inphase current vs antiphase current | N = 7; Rank sum = 28 | 0.017 | X |
| Fig 5E | Unpaired Wilcoxon rank sum, one-tailed | Ctrl vs. KO Thal dist 0-0.5 mm | ranksum: 2249, zstat: 6.5971 | 2.10e-11 | X |
|  | Unpaired Wilcoxon rank sum, one-tailed | Ctrl vs. KO Thal dist 0.5-1 mm | ranksum: 6432, zstat: 5.9760 | 1.14e-09 | X |
|  | Unpaired Wilcoxon rank sum, one-tailed | Ctrl vs. KO Thal dist 1-1.5 mm | ranksum: 1318, zstat: 4.3054 | 8.33e-06 | X |
|  | Unpaired Wilcoxon rank sum, one-tailed | Ctrl vs. KO Ctx dist 2-2.5 mm | ranksum: 513, zstat: 4.0381 | 2.69e-05 | X |
|  | Unpaired Wilcoxon rank sum, one-tailed | Ctrl vs. KO Ctx dist 2.5-3 mm | ranksum: 3114, zstat: 4.3320 | 7.39e-06 | X |
| Fig 5H (histograms) | Unpaired Wilcoxon rank sum, two-tailed | DN Ctrl Still vs. DN KO Still | ranksum: 9694998, zstat: -6.3873, Cohen’s *d*: -0.1909 | 1.69e-10 | X |
|  | Unpaired Wilcoxon rank sum, one-tailed | Thal Ctrl Still vs. Thal KO Still | ranksum: 12209905, zval: 34.3450, Cohen’s *d*: 1.0114 | 8.35e-259 | X |
|  | Unpaired Wilcoxon rank sum, one-tailed | Ctx Ctrl Still vs. Ctx KO Still | ranksum: 11328214, zval: 20.0648, Cohen’s *d*: 0.5692 | 7.49e-90 | X |
| Supp Fig 7J (histograms) | Unpaired Wilcoxon rank sum, two-tailed | DN Ctrl Move vs. DN KO Move | ranksum: 1567951, zstat: -0.4148, Cohen’s d: -0.0055 | 0.6783 |  |
|  | Unpaired Wilcoxon rank sum, two-tailed | Ctx Ctrl Move vs. Ctx KO Move | ranksum: 1649748, zstat: 4.8670, Cohen’s d: 0.1738 | 1.13e-06 | X |
|  | Unpaired Wilcoxon rank sum, two-tailed | Thal Ctrl Move vs. Thal KO Move | ranksum: 1489911, zstat: -5.4541, Cohen’s d: -0.2048 | 1 |  |
| Fig 6B | Kruskal-Wallis, FDR corrected (Benjamini-Hochberg) | Ctrl vs. KO Step Count | 0.0533 | 0.9795 |  |
|  |  | Ctrl vs. KO Avg Velocity | 3.3143 | 0.4121 |  |
|  |  | Ctrl vs. KO Step Width – LF Paw | 3.8981 | 0.4121 |  |
|  |  | Ctrl vs. KO Step Width – LH Paw | 1.7101 | 0.5712 |  |
|  |  | Ctrl vs. KO Step Width – RF Paw | 4.1032 | 0.4121 |  |
|  |  | Ctrl vs. KO Step Width – RH Paw | 1.2156 | 0.5712 |  |
|  |  | Ctrl vs. KO Stride Length – LF Paw | 1.1052 | 0.5712 |  |
|  |  | Ctrl vs. KO Stride Length – LH Paw | 1.3314 | 0.5712 |  |
|  |  | Ctrl vs. KO Stride Length – RF Paw | 1 | 0.5712 |  |
|  |  | Ctrl vs. KO Stride Length – RH Paw | 1.2156 | 0.5712 |  |
| Fig 6C | Kruskal-Wallis, FDR corrected (Benjamini-Hochberg) | Ctrl vs. KO Duty Factor – Front | 0.1111 | 0.9795 |  |
|  |  | Ctrl vs. KO Duty Factor – Hind | 0.0007 | 0.9795 |  |
|  |  | Ctrl vs. KO DF Imbalance – Front | 0.0532 | 0.9795 |  |
|  |  | Ctrl vs. KO DF Imbalance – Hind | 1.8468 | 0.9795 |  |
|  |  | Ctrl vs. KO Temporal Symmetry – Front | 0.0007 | 0.9795 |  |
|  |  | Ctrl vs. KO Temporal Symmetry – Hind | 0.0164 | 0.5712 |  |
|  |  | Ctrl vs. KO Spatial Symmetry – Front | 0.0059 | 0.9795 |  |
|  |  | Ctrl vs. KO Spatial Symmetry – Hind | 0.0796 | 0.9795 |  |
| Fig 6D | 3-way ANOVA | Training Day | F-stat: 1.45 | 0.1643 |  |
|  |  | Trial (first vs. final) | F-stat: 22.32 | 0 | X |
|  |  | Group (Ctrl vs. KO) | F-stat: 39.86 | 0 | X |
|  |  | Trial:Training Day | F-stat: 0.12 | 0.9991 |  |
|  |  | Trial:Group | F-stat: 7.68 | 0.0058 | X |
|  |  | Training Day:Group | F-stat: 0.61 | 0.7913 |  |
| Figure 6E | 2-way ANOVA | Training Day | F-stat: 28.35 | 0 | X |
|  |  | Group (Ctrl vs. KO) | F-stat: 62.43 | 0 | X |
|  |  | Training Day:Group | F-stat: 0.6 | 0.7971 |  |
| Figure 6F | Unpaired Wilcoxon rank sum, two-tailed | Intra-day improvement | ranksum: 209.5, zstat: 2.8834 | 0.0039 | X |
| Supp Fig 8C-D | Kruskal-Wallis, FDR corrected (Benjamini-Hochberg) | Ctrl vs. KO Distance Traveled (pixels) | 3.8981 | 0.1731 |  |
|  |  | Ctrl vs. KO Avg Pawprint Size | 0.0532 | 0.8530 |  |
|  |  | Ctrl vs. KO L/R Paw Pressure Index Ratio – Rearing | 0.3793 | 0.6623 |  |
|  |  | Ctrl vs. KO L/R Paw Pressure Index Ratio – Standing | 0.8532 | 0.5336 |  |
|  |  | Ctrl vs. KO F/H Paw Pressure Index Ratio | 7.1331 | 0.0454 | X |
|  |  | Ctrl vs. KO Hind Paw Lifted Time – Left | 0.0796 | 0.8530 |  |
|  |  | Ctrl vs. KO Hind Paw Lifted Time – Right | 0.4109 | 0.6624 |  |
|  |  | Ctrl vs. KO Rearing Time | 2.9513 | 0.2574 |  |

**Supplemental Data Table 1. Statistics**
